# Therapeutic ability of extracellular vesicles derived from angio-miRNA-modified mesenchymal stromal cells in ischemic animal models

**DOI:** 10.1101/2025.01.27.635184

**Authors:** Yoshiki Wada, Toshifumi Kudo, Tomomi Kusakabe, Ayako Inoue, Yusuke Yoshioka, Takahiro Ochiya, Shoji Fukuda

## Abstract

**Background:** Regenerative therapy using extracellular vesicles (EVs) is expected to be an option for supportive therapy of chronic limb-threatening ischemia. In this study, we examined the angiogenic potential of EVs derived from genetically modified MSCs, focusing on the angio-miRNAs in these vesicles.

**Methods:** Bone marrow mesenchymal stromal cells (BM-MSCs) were transfected with a lentiviral vector containing specific angio-miRNAs (miRNA-126, -135b, or -210). EVs were separated from the culture medium of BM-MSCs via ultracentrifugation. The quality and quantity of the EVs were evaluated using nanoparticle tracking analysis, fluorometry, and an ExoScreen assay. Tube formation assays were conducted using human umbilical vein endothelial cells to assess the angiogenic potency of the EVs *in vitro*. The EVs were injected into the ischemic hind limb muscles of mice. The grades of limb ischemia, blood perfusion, and muscle tissues were analyzed to assess the angiogenic potency of the EVs *in vivo*.

**Results:** Angio-miRNAs were overexpressed in MSCs transfected with a lentiviral vector. Tube formation was significantly enhanced when endothelial cells were cultured with the EVs (P < 0.05). Compared to non-contact co-culture with BM-MSCs, EVs derived from modified BM-MSCs significantly promoted tube formation in endothelial cells (P < 0.05). The severity of limb ischemia on day 7 was alleviated when a combination of two or three EVs derived from modified BM-MSCs was injected compared with native EVs. Toe blood perfusion on day 14 significantly recovered when a combination of two or three EVs derived from modified BM-MSCs was injected (P < 0.05). Tissue analysis revealed that the number of newly formed endothelial cells in ischemic muscles increased when a combination of two or three EVs derived from modified BM-MSCs was injected.

**Conclusions:** The use of a combination of EVs derived from genetically modified MSCs stimulated angiogenesis both *in vitro* and *in vivo*.

## Introduction

Chronic limb-threatening ischemia (CLTI) has a one-year lower limb amputation rate of 22% and a mortality rate of 22% when revascularization is unsuccessful or inapplicable^1^. These outcomes have serious adverse effects on activities of daily living and life expectancy^2^. Regenerative therapy for CLTI has been attempted using stem cells^3^, mononuclear cells^4^, and growth factors^5^. Fukuda et al. implemented angiogenic therapy using platelet-rich plasma and bone marrow-derived mesenchymal stromal cells (BM-MSCs)^6^. The paracrine effect may contribute to treating lower limb ischemic ulcers with MSCs^7^.

Although MSCs were initially expected to be used for cell therapy because of their pluripotency, recent research has suggested that cytokines, growth factors, and microRNAs (miRNAs) secreted from these cells may also have therapeutic effects^8^. Cells secrete cargo by packaging them in extracellular vesicles (EVs)^9^. Therefore, the potential therapeutic applications of MSC-derived EVs have been considered. Proteins, DNAs, long non-coding RNAs, mRNAs, and miRNAs contained within EVs can be released outside the cells, altering gene expression and inducing angiogenesis in the surrounding cells^10^.

MSC-derived EVs contain miRNAs that induce angiogenesis, known as angio-miRNAs^11^. In this study, we focused on three angio-miRNAs, miRNA-126, -135b, and -210. This study aimed to investigate whether EVs combined with angio-miRNAs could promote angiogenesis *in vitro* and *in vivo* to search for clinically applicable regenerative therapy with EVs for CLTI patients.

## Methods

### Culture of BM-MSCs

Human BM-MSCs were obtained from Lonza (lot no. TL281098). The cells were seeded at a density of 2.0 × 10^5^ per 10-cm dish in 8 ml of MesenPro (Thermo Fisher Scientific, USA) medium supplemented with 1% GlutaMAX^TM^ (Thermo Fisher Scientific, USA) and incubated at 37℃ and 5% CO_2_. BM-MSCs from the fourth passage were used for the experiments.

### Generation of BM-MSCs overexpressing angio-miRNAs

A lentiviral system was used to overexpress miRNA-126, -135b, and -210 in MSCs. Briefly, lentiviral stocks containing human miRNA precursors were purchased from BioSettia Inc, USA. MSCs were plated on 6-well plates at a density of 1.0 × 10^5^ cells and incubated for 24 h. Afterward, the spent medium was replaced with a medium containing TransDux™ (System Biosciences, USA), and MSCs were incubated with the lentiviral particles for 24 h. MSCs in the lentiviral vector group were transfected with lentiviral vectors containing miRNA-126, -135b, and -210. Each vector was transfected at a multiplicity of infection (MOI) of 1. Similarly, MSCs in the control vector group were infected with a control vector (a negative control lentiviral vector containing a scrambled sequence) at an MOI of 1. The cell pools were then selected using a medium containing 2.0 μg/ml puromycin for 3 days after transfection. The genetically modified cells were named MSC-miRNA126, MSC-miRNA135b, MSC-miRNA210, and MSC-miRNA controls.

### MiRNA extraction and qPCR

Total miRNA from each MSC culture was extracted using the miRNeasy Kit for miRNA Purification (Qiagen, Netherlands) following the manufacturer’s protocol. cDNA synthesis was performed using a TaqMan™ MicroRNA Reverse Transcription Kit (Thermo Fisher Scientific, USA). Real-time qPCR was performed using primers specific for miRNA-126, - 135b, and -210. U6 was used as an internal control.

### Separation and characterization of EVs

Separation of EVs was conducted according to the following procedure. Briefly, 7 × 10^5^ BM-MSCs were passaged in a 15-cm dish containing 20 ml of StemPro (Thermo Fisher Scientific, USA). After 24 h, the BM-MSC culture medium was collected and centrifuged at 2,000 ×*g* for 10 min. The supernatant was then filtered through a 0.22-μm pore membrane filter and poured into 13.2-ml Open-Top Thinwall Ultra-Clear Tubes (Beckman Coulter, USA). The tubes were ultracentrifuged using an SW 41 Ti rotor (Beckman Coulter, USA) at 35,000 rpm at 4℃ for 70 min. The supernatant was decanted, and the precipitated EVs were diluted with phosphate-buffered saline (PBS(–)). The EV solution was then ultracentrifuged under the same conditions, the supernatant was decanted, and the pellets were collected. Each type of EV isolated was named after the vector transfected into its BM-MSC source (EVcontrol, EV126, EV135b, or EV210); EVs derived from native BM-MSCs were named EVNative. The number of EVs was measured by determining the concentrations of EV-associated proteins using a Qubit Fluorometer (Thermo Fisher Scientific, USA). The quality of the EVs was verified through dynamic light scattering using a NanoSight instrument (Spectris, UK) following the manufacturer’s instructions. The ExoScreen assay was performed as previously described^12^ to capture CD63, an EV surface marker. Briefly, 96-well half-area white plates (6002290; Revvity, USA) were filled with 10 μl of EVs samples and 15 μl of a mixture of anti-human CD63 antibody solid-phase acceptor beads (6772002; Revvity, USA) and biotinylated anti-human CD63 antibodies. The reaction mixture was incubated at 37℃ for 90 min in the dark. Without a washing step, 25 µl of AlphaScreen streptavidin-coated donor beads (6760002; Revvity, USA) were added to the mixture. The reaction mixture was then incubated in the dark for another 45 min at 37℃. Finally, the absorbance of each well was detected on an EnSight Multilabel Plate Reader (Revvity, USA) at an excitation wavelength of 680 nm and emission wavelength of 615 nm. Background signals from 0.22-μm filtered PBS(–) were subtracted from the measured signals. The above procedures were performed following the minimal information available for studies on EVs (MISEV2023)^13^.

### Tube formation assay

This study was conducted in accordance with the Tokyo Medical University Medical Ethics Committee and the Regulations Concerning Medical Research(D23-036). The human-derived cells used in this study were obtained from ethically approved sources (Lonza, Switzerland) and complied with the supplier’s regulations and the Declaration of Helsinki.

Human umbilical vein endothelial cells (HUVECs) (Lonza, Switzerland) were seeded at a density of 2.0×10^5^ cells per 10-cm dish in 8 ml of EBM-2 medium (Lonza, Switzerland) and incubated at 37℃ and 5% CO_2_. Cells from the fifth to ninth passages were used in the tube formation assay.

HUVECs and EVs from BM-MSCs (0.5, 1, or 10 μg/ml) suspended in EBM-2 medium were seeded on top of a Matrigel (Corning, USA) in a 24-well plate at a density of 1.5 × 10^4^ cells and incubated at 37°C for 24 h. In the control group, HUVECs were seeded onto Matrigel in PBS(–). Images were captured using a BZ-X 800 phase contrast microscope (Keyence, Japan), and tube formation was analyzed using the Angiogenesis Analyzer in ImageJ software (version 1.54d; https://imagej.net/; National Institutes of Health). The number of junctions, total length of segments, and total mesh areas counted by the software were compared between each concentration and the control group.

We also performed tube formation analysis of HUVECs cultured with single EVs or combined EVs (1 μg/ml) derived from transfected BM-MSCs. Combined EVs refer to a mixture of two or three EVs (EV126 and EV135b; EV135b and EV210; EV126 and EV210; and a combination of EV126, EV135b, and EV210). To analyze the angiogenic capacity of the BM-MSC culture medium, a non-contact co-culture model was employed by placing a Transwell insert (Corning, USA) on each well. Briefly, 1.0 × 10^4^ BM-MSCs were seeded on Transwell plates containing MesenPro medium. Tube formation by HUVECs was analyzed after they were cultured separately from BM-MSCs in Transwell plates for 24 h. The number of junctions, total length of segments, and total mesh areas counted by the software were compared between native EVs, transfected EVs, and co-culture models.

### Mouse hindlimb ischemia model

Animal studies were conducted in accordance with the National Institutes of Health Guide for the Care and Use of Laboratory Animals. The animal study protocols were approved by the Institutional Animal Care and Use Committee of Tokyo Medical University (R6-020). Briefly, 10-week-old male BALB/c mice were purchased from Japan SLC Inc. (Japan). The mice were allocated to 11 groups: sham surgery, PBS(–), native EVs, control EVs, single EVs (EV126, EV135b, or EV210), double EVs (EV126 and EV135b, EV135b and EV210, and EV126 and EV210), and triple EVs (EV126, EV135b, and EV210) (n=4 mice for each group). Mice were anesthetized via intraperitoneal injection of a mixture of midazolam (0.4 mg/ml), butorphanol (0.5 mg/ml), and medetomidine (0.075 mg/ml). After sedation, a hair removal cream was applied bilaterally to the abdomen and hind limbs. Both hind limbs were fixed with extension and abduction in the supine position. Skin without hair was cleansed with a 80% ethanol swab, and an oblique incision was made on the left groin using surgical scissors. The subcutaneous fat pad was then dissected using a cotton swab. The femoral artery, vein, and nerve are identified and gently separated. The common femoral artery was ligated at two points using a 6-0 polyvinylidene fluoride suture (Kono Seisakusho, Japan). Unilateral hindlimb ischemia was induced by excising the left common femoral artery between the two ligations. After the excision of the common femoral artery, 50 μl of EVs containing 4 μg protein that were measured by Qubit Fluorometry were injected into the extensor and flexor muscles of the thigh from the surgical site using 1 ml syringes with 27-gauge needles (Terumo, Japan). PBS(–) was injected into the rats in the control group. The incision was closed using 4-0 silk sutures (Kono Seisakusho, Japan), and the surgical wound was sterilized using povidone-iodine. The right hind limb was used as an internal control. The mice were laid on a heating pad kept at 37℃ to avoid body temperature reduction. Four mice underwent a sham operation in which the left common femoral artery was exposed but not ligated.

### Evaluation of limbs

The severity of ischemic injury was evaluated based on the grade of limb necrosis, according to a previous study^14^: grade 0, normal limb without necrosis; grade I, black toenails with necrosis limited to the toes; grade II, necrosis extending to the foot; grade III, necrosis extending to the knee; and grade IV, necrosis extending to the hip or loss of the entire limb.

Blood flow was monitored using a MoorFLPI-2 instrument (Moor Instruments, UK) on days 0 (before and after the procedure) and 7. To evaluate alterations in limb perfusion, the blood flow ratio of the left foot to the right foot was analyzed. The region of interest was set at the ankle, which reflected peripheral blood perfusion.

### Histological and immunohistochemical analyses

Limb muscle tissues were immunohistochemically stained with hematoxylin-eosin and anti-CD31 antibodies (GeneTex International Corporation, USA). Briefly, the left gastrocnemius muscle was harvested from mice in each group on day 14 (n=1 per group) and fixed with a 10% formalin solution (Muto Pure Chemicals, Japan). The samples were then embedded in paraffin, and 3-μm-thick serial sections were processed for hematoxylin-eosin staining.

For immunostaining to detect CD31, sections were incubated with anti-CD31 antibodies (1:4000) overnight at 4℃ and subsequently incubated with the secondary antibody, histofine simple stain mouse MAX-PO^®^ (1:1; Nichirei, Japan) for 30 min at room temperature. All sections were observed under a BZ-X 800 microscope (Keyence, Japan).

To evaluate angiogenesis in the ischemic muscle, we compared the ratio of the area occupied by capillary endothelial cells in the ischemic muscle between mice injected with EVs and PBS(–). The areas of muscle and capillary endothelial cells were calculated using ImageJ software.

### Statistics

The results were expressed as the mean ± standard error of the mean. Differences between groups were determined using analysis of variance (ANOVA). All data were analyzed using EZR (Saitama Medical Center, Jichi Medical University, Saitama, Japan), a graphical user interface for R (R Foundation for Statistical Computing, Vienna, Austria). Statistical significance was set at P < 0.05.

## Results

### Establishment of transfected MSCs

Transfection of miRNAs in MSCs was confirmed via red fluorescent protein (RFP) expression, which was visualized using fluorescence microscopy (Figure 1). Table 1 shows the qPCR results. Relative to that of U6, the fold change in the expression of miRNA-126 in MSC-miRNA126 was 395.2, that of miRNA-135b in MSC-miRNA135b was 132.4, and that of miRNA-210 in MSC-miRNA210 was 8.84. However, the fold change in the MSC-miRNA control did not change (0.8584 for miRNA-126, 1.04 for miRNA-135b, and 1.082 for miRNA-210) compared to those of the internal control. These results indicate that the expression of transfected miRNAs in each MSC type was elevated.

**Figure 1.**
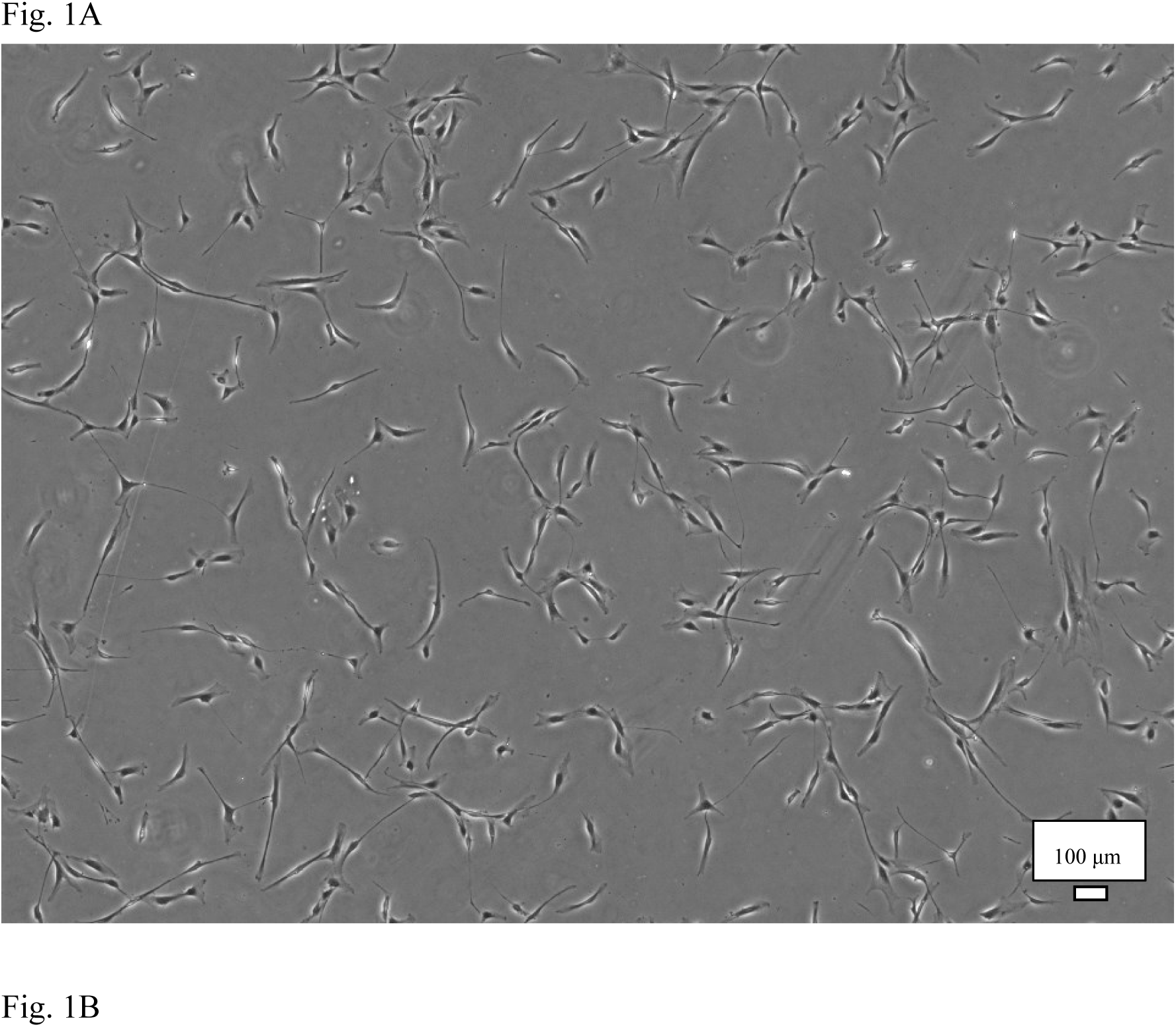
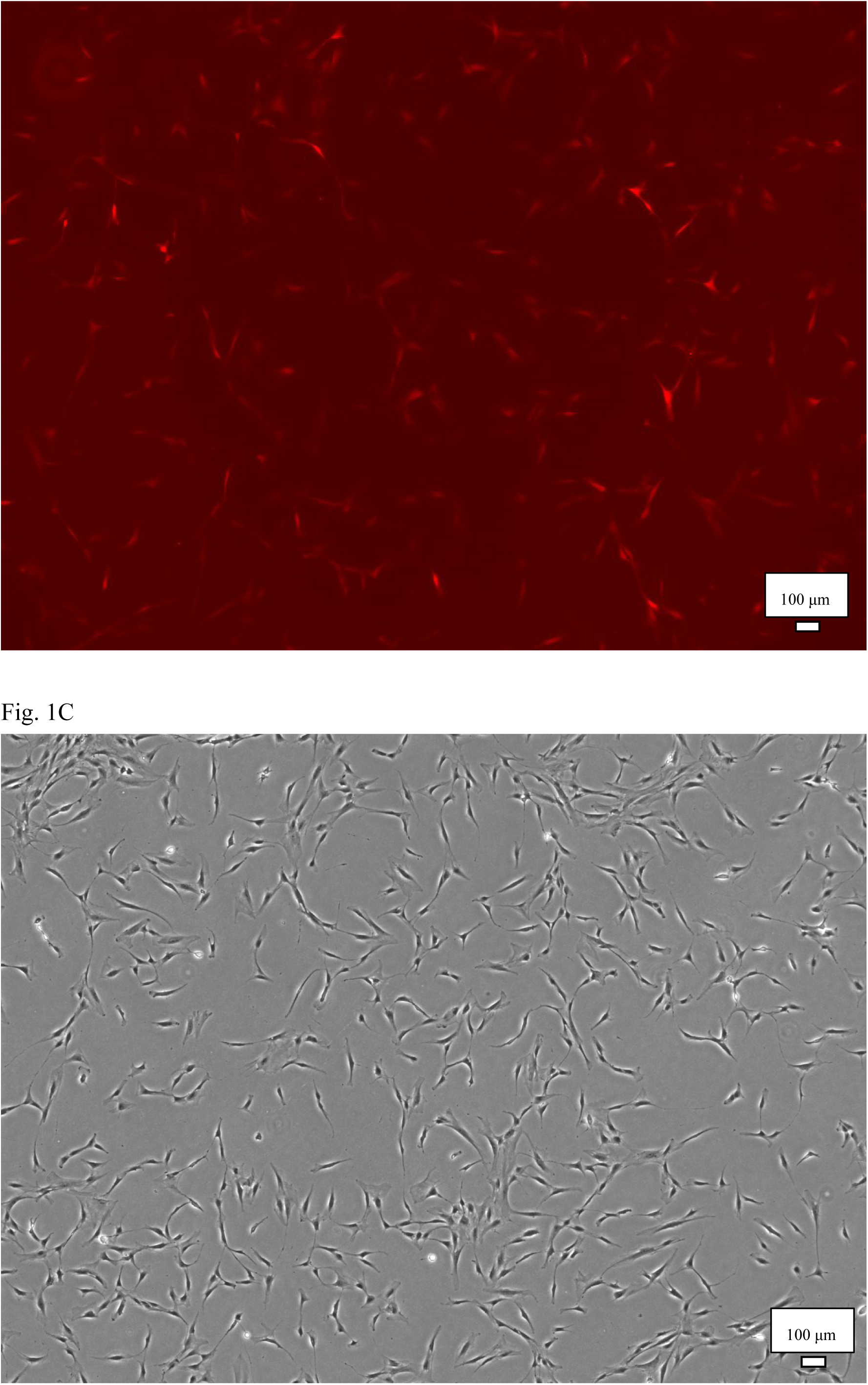
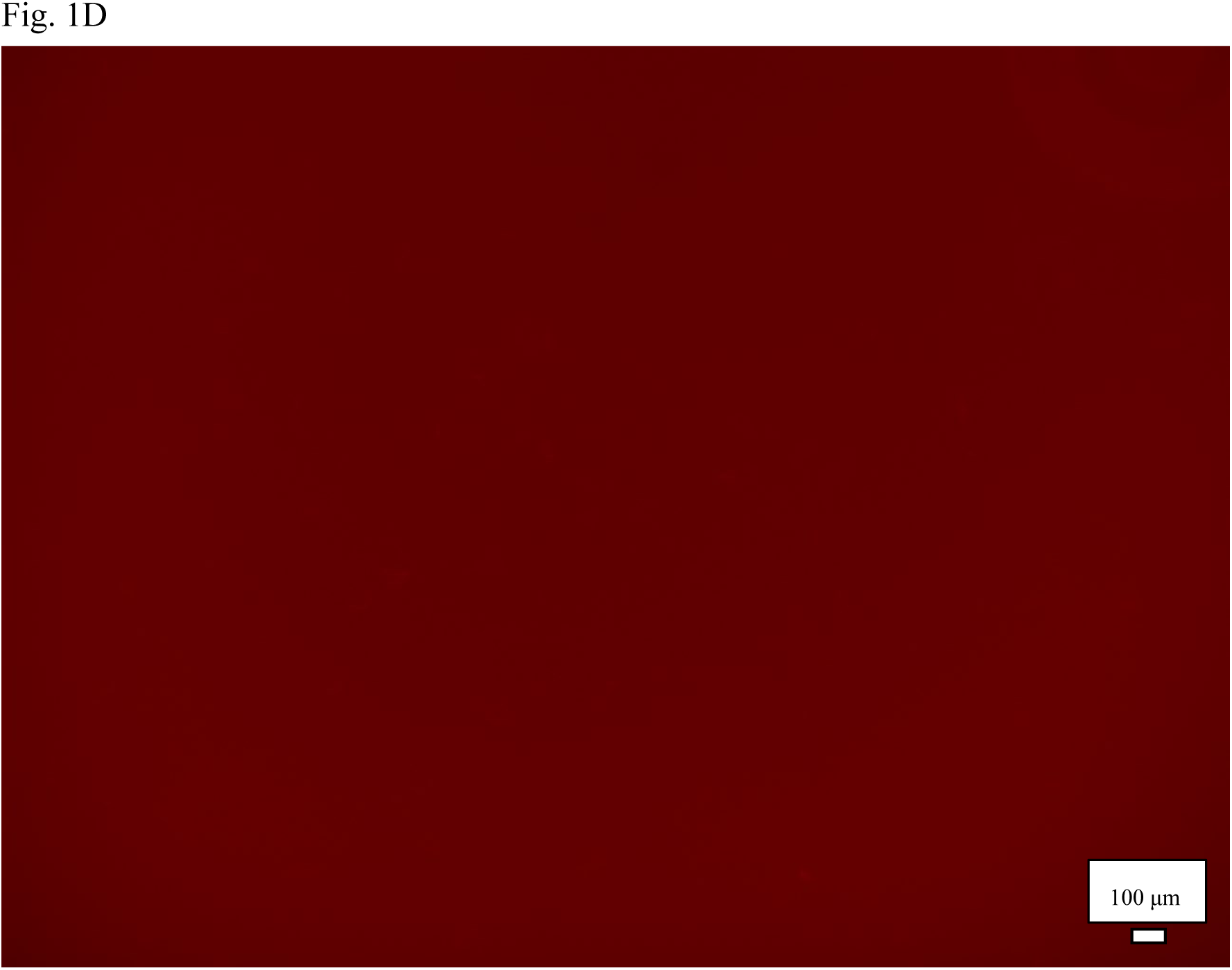
Transfection of bone marrow-derived mesenchymal stromal cells (BM-MSCs). A, BM-MSCs transduced with the lentiviral vector. B, Red fluorescent protein (RFP) fluorescence proved that the vector was integrated into the BM-MSC genome. C and D, native BM-MSCs do not express RFP.

**Table 1.** Relative gene expression of angio-miRNAs in modified mesenchymal stromal cells (MSCs) compared with native MSCs.

| Fold change (relative<br>to U6 expression) | miRNA-126 | miRNA-135b | miRNA-210 |
| --- | --- | --- | --- |
| MSC-miR control | 0.8584 | 1.041 | 1.082 |
| MSC-miR126 | 395.2 | 0.8424 | 1.214 |
| MSC-miR135b | 0.7409 | 132.4 | 1.137 |
| MSC-miR210 | 0.1828 | 0.443 | 8.842 |

### Characterization of EVs

The volume of the culture medium and the total cell number used to separate each EV are presented in Table 2. The protein concentrations of each EV are listed in Table 3. The particle concentrations and size distributions are presented in Table 4 and Figure 2, respectively. The protein-to-particle ratios are presented in Table 5. Specifically, the protein-to-particle ratios of EVNative, EVcontrol, EV126, EV135b, and EV210 were 1.71 × 10^-8^, 1.68 × 10^-8^, 8.74 × 10^-9^, 1.20 × 10^-8^, and 1.31 × 10^-8^, respectively. ExoScreen analysis confirmed the presence of CD63 in all samples (Figure. 3). These findings indicate that the quality of EVs derived from BM-MSCs aligned with the MISEV2023 standards.

**Figure 2.**
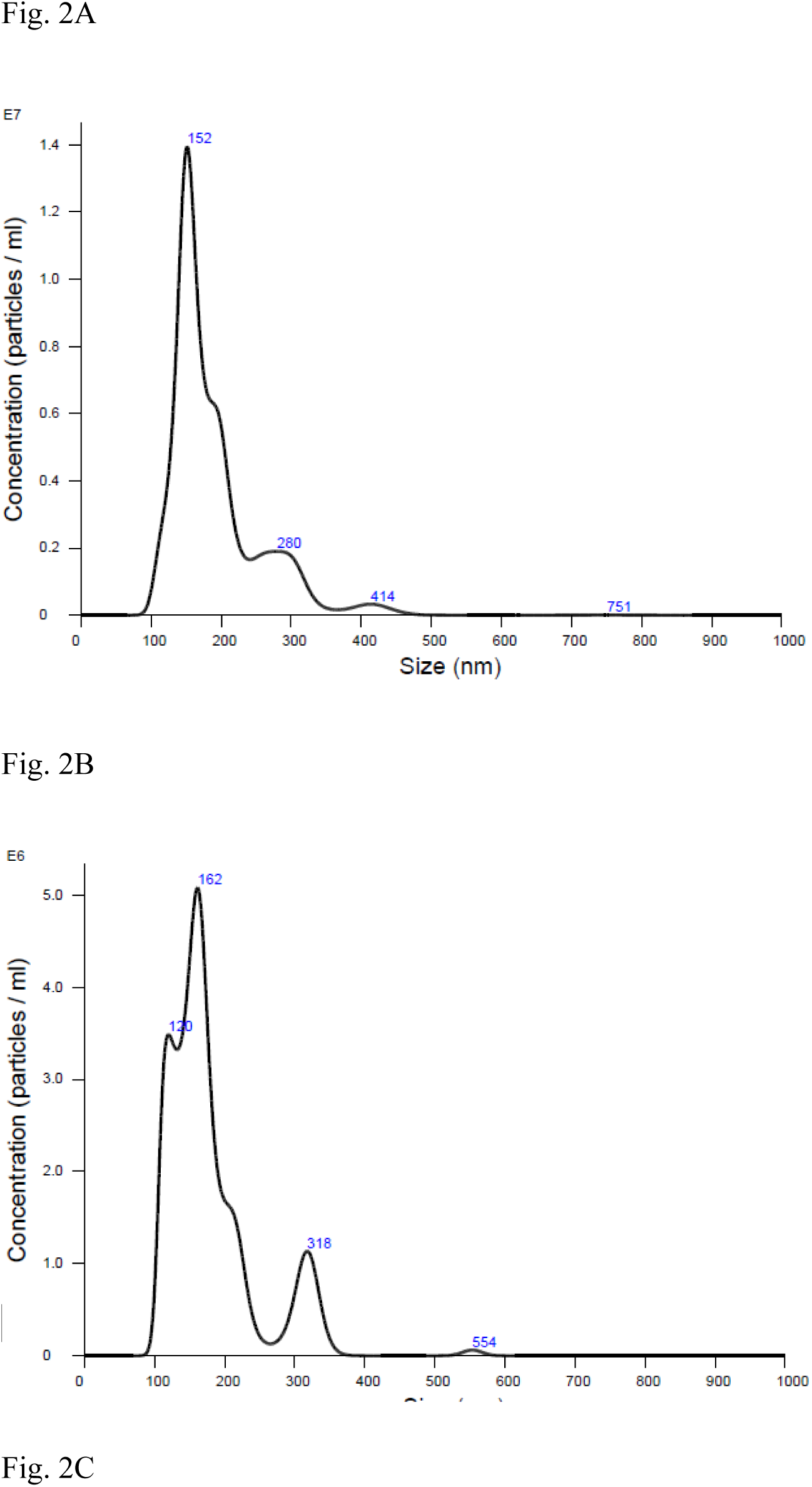
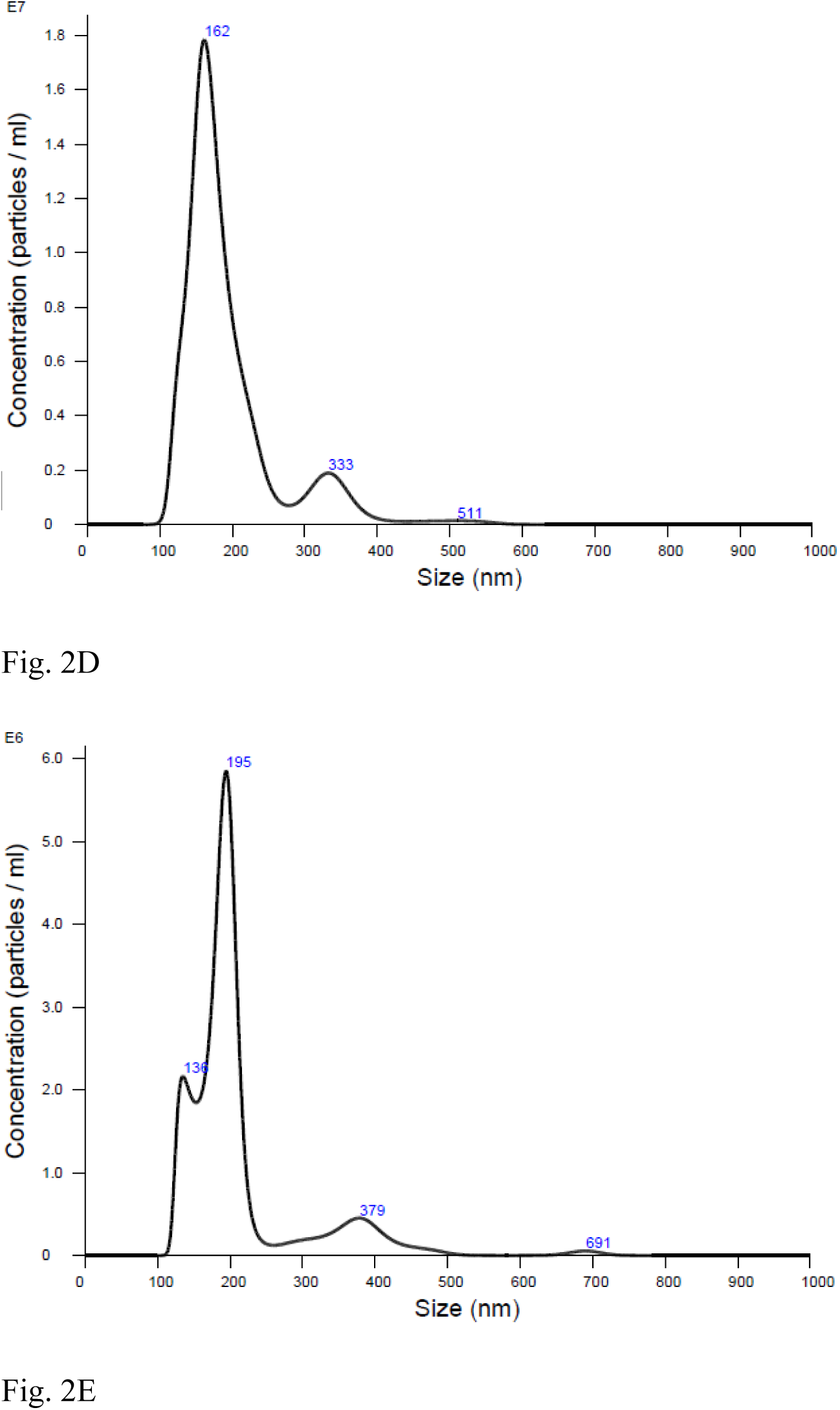
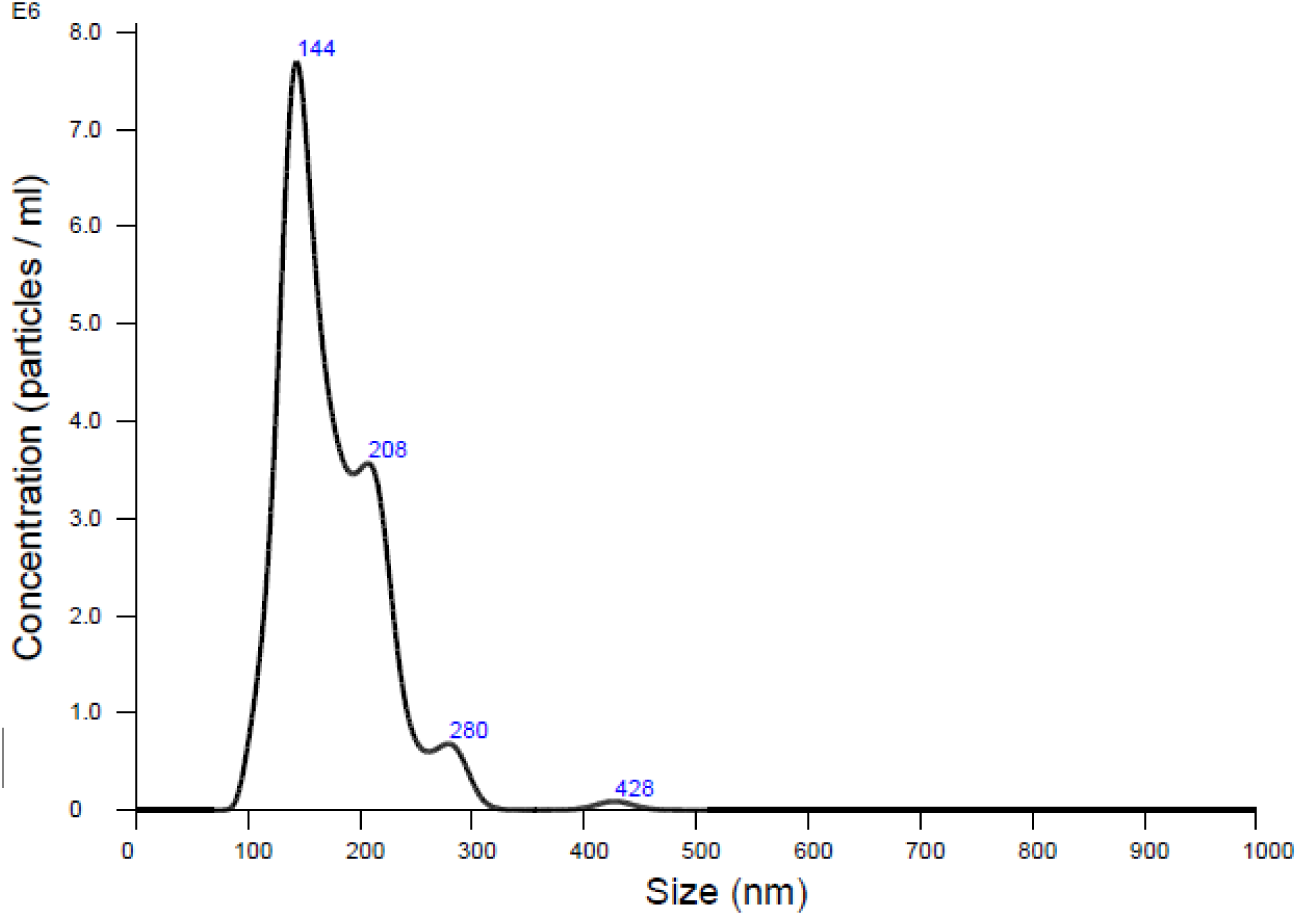
Particle size distribution in extracellular vesicle (EV) samples derived from each type of modified mesenchymal stromal cell. A, EVNative. B, EVcontrol. C, EV126. D, EV135b. E, EV210.

**Figure 3.**
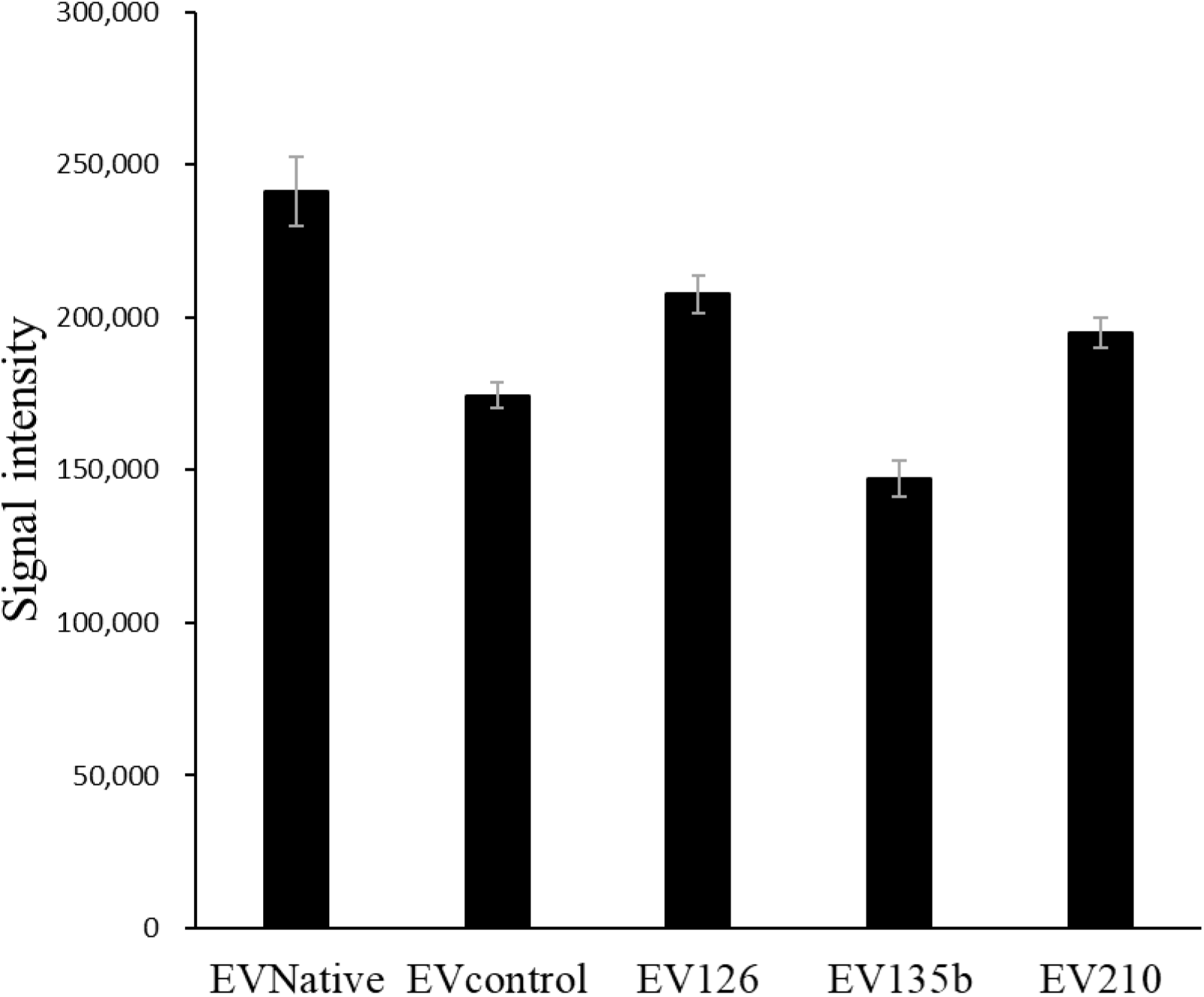
The result of ExoScreen assay. Data are presented as the mean ± standard error of the mean. (N = 3 samples for each group).

**Table 2.** Culture medium and total cell number used for extracellular vesicle separation.

|  | Culture medium (ml) | Total cell number |
| --- | --- | --- |
| EVNative | 272.6 | $9.17 \times 10^6$ |
| EVcontrol | 179.4 | $6.50 \times 10^6$ |
| EV126 | 220.8 | $9.22 \times 10^6$ |
| EV135b | 218.2 | $0.88 \times 10^6$ |
| EV210 | 186.4 | $8.60 \times 10^6$ |

**Table 3.** Protein concentration as measured using a fluorometer.

|  | Protein concentration (µg/ml) |
| --- | --- |
| EVNative | 1714 |
| EVcontrol | 742 |
| EV126 | 1216 |
| EV135b | 440 |
| EV210 | 784 |

**Table 4.** Particle size distribution and concentration as measured using NanoSight.

|  | mean, nm | mode, nm | SD, nm | D10, nm | D50, nm | D90, nm | Concentration,<br>particles/ml |
| --- | --- | --- | --- | --- | --- | --- | --- |
| EVNative | 189 | 151.6 | 63.4 | 132.5 | 168.7 | 282.9 | $1.02 \times 10^{11}$ |
| EVcontrol | 179.7 | 161.3 | 65.3 | 117.7 | 162.9 | 299 | $4.41 \times 10^{10}$ |
| EV126 | 193.4 | 161.4 | 68.1 | 135.7 | 173.2 | 301.2 | $1.39 \times 10^{11}$ |
| EV135b | 212.4 | 194.6 | 85.9 | 140.6 | 191.6 | 353 | $3.67 \times 10^{10}$ |
| EV210 | 174.1 | 143.5 | 46.8 | 126.4 | 163.1 | 230.4 | $6.0 \times 10^{10}$ |

**Table 5.** Protein-to-particle ratio of each type of extracellular vesicle.

|  | Protein/particles (μg/particle) |
| --- | --- |
| EVNative | $1.71 \times 10^{-8}$ |
| EVcontrol | $1.68 \times 10^{-8}$ |
| EV126 | $8.74 \times 10^{-9}$ |
| EV135b | $1.20 \times 10^{-8}$ |
| EV210 | $1.31 \times 10^{-8}$ |

### EVs induced tube formation in HUVECs

Tube formation in HUVECs cultured with and without EVNative was analyzed using ImageJ software (Figure 4A-4C). The results showed a significant increase in tube formation parameters when HUVECs were cultured with EVNative (P < 0.05) (Figure 4D). We also compared the tube formation ability of HUVECs cultured with single EVs, combined EVs, and BM-MSCs (Figure 5A-5J). The results showed that EV126, EV135b, and EV126+EV135b significantly enhanced the three tube formation parameters compared to co-culture with BM-MSCs alone (P < 0.05). Moreover, EVs derived from transfected BM-MSCs tended to increase tube formation compared to native BM-MSCs, although this was not statistically significant (Figure 5K).

**Figure 4.**
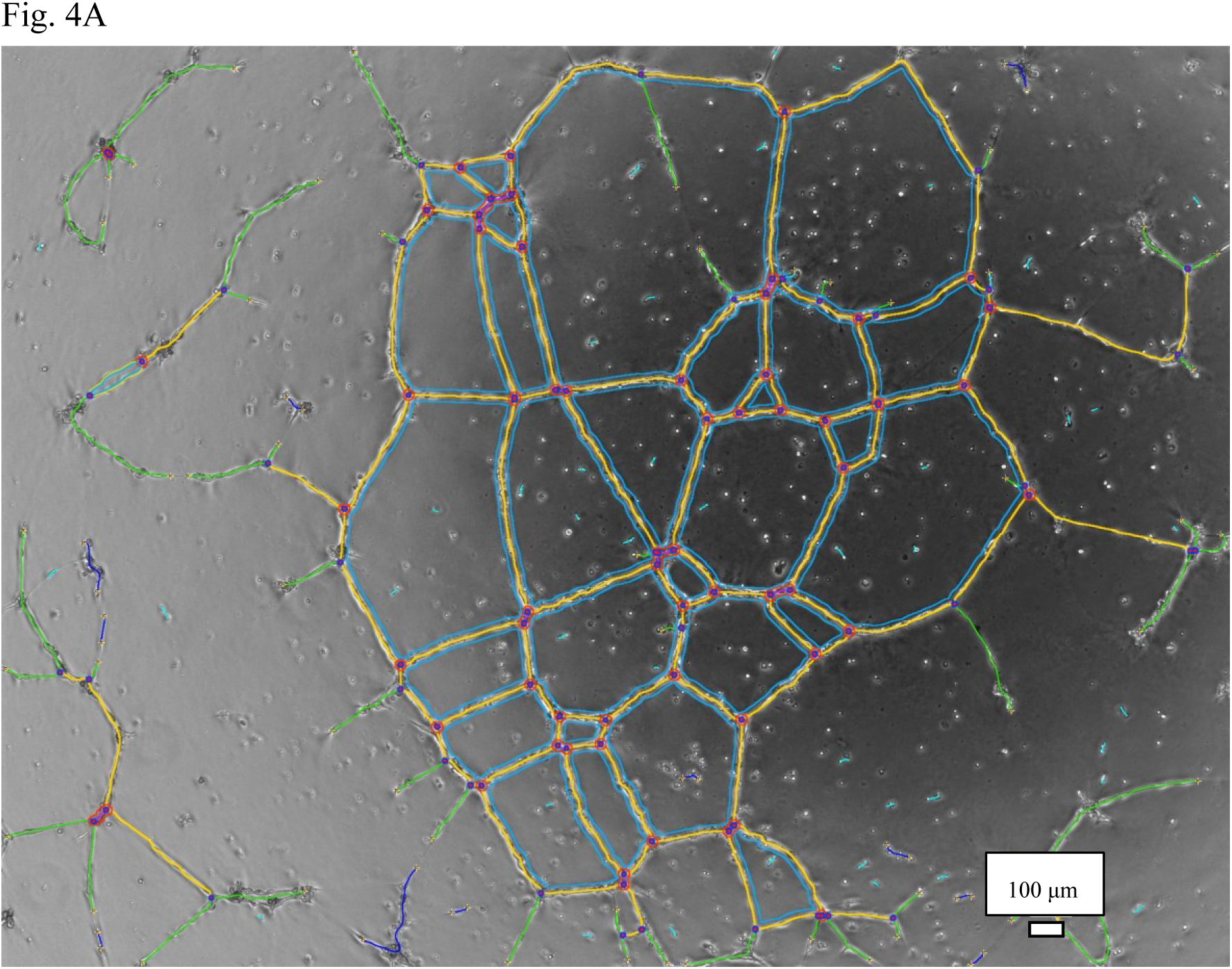
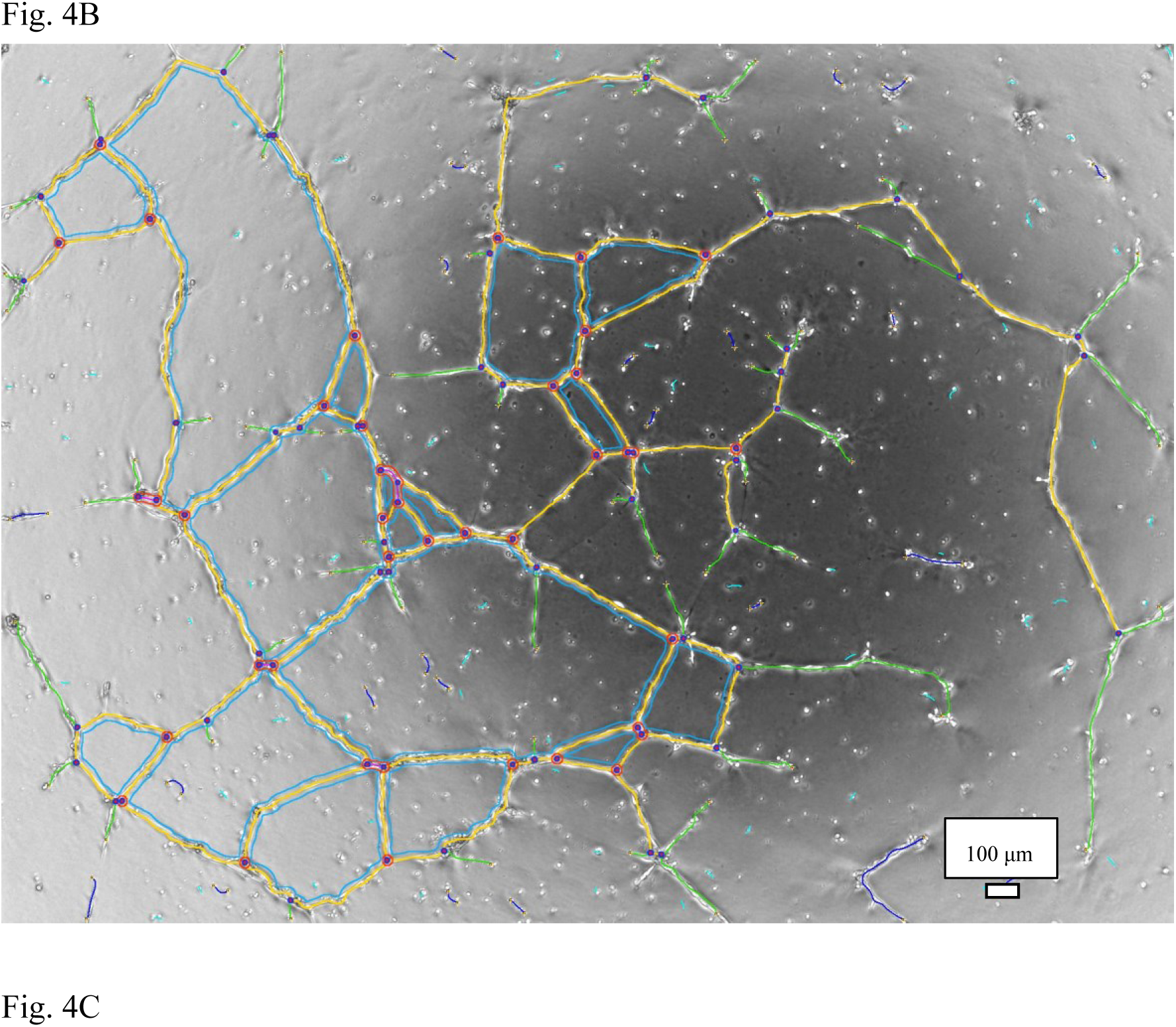
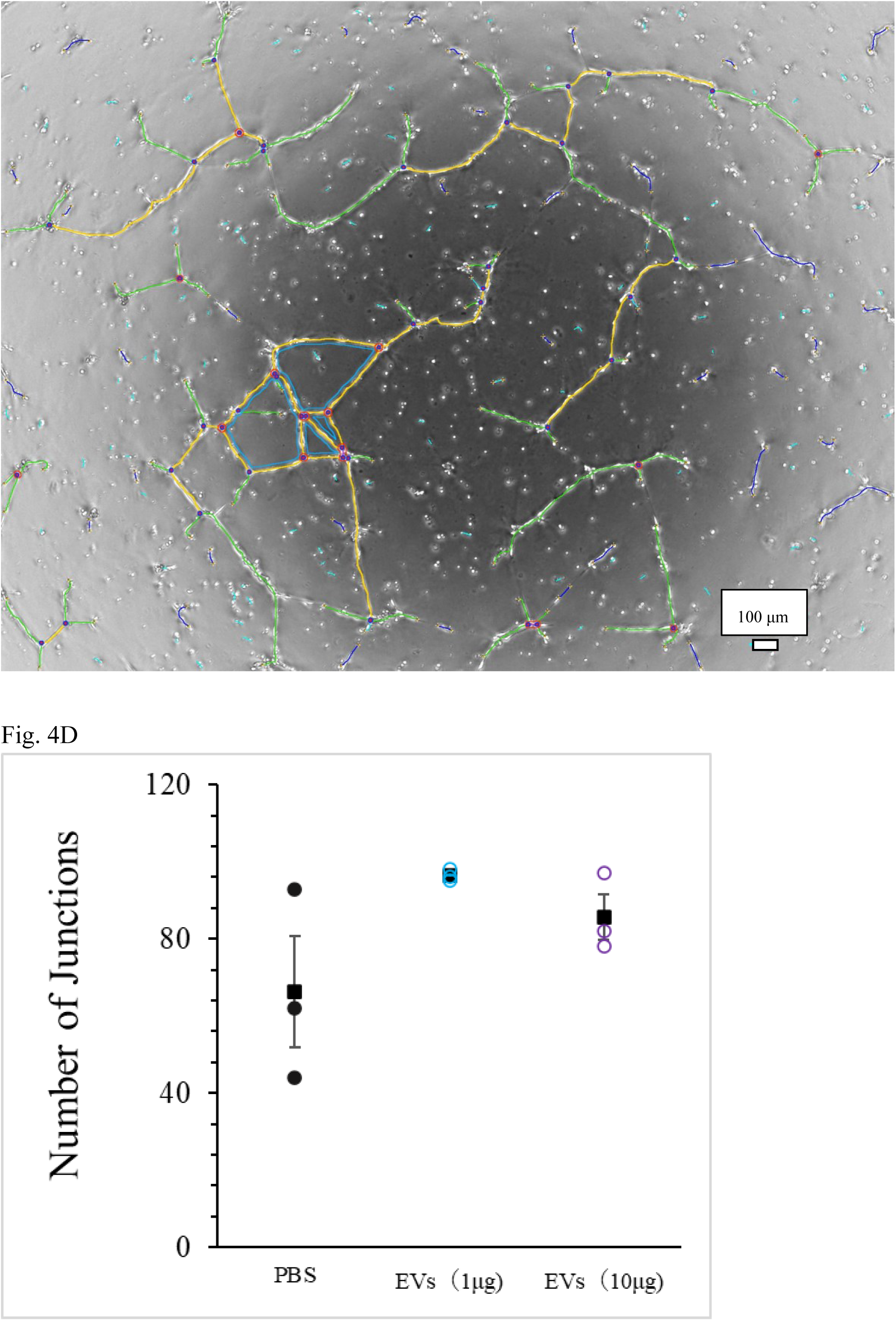
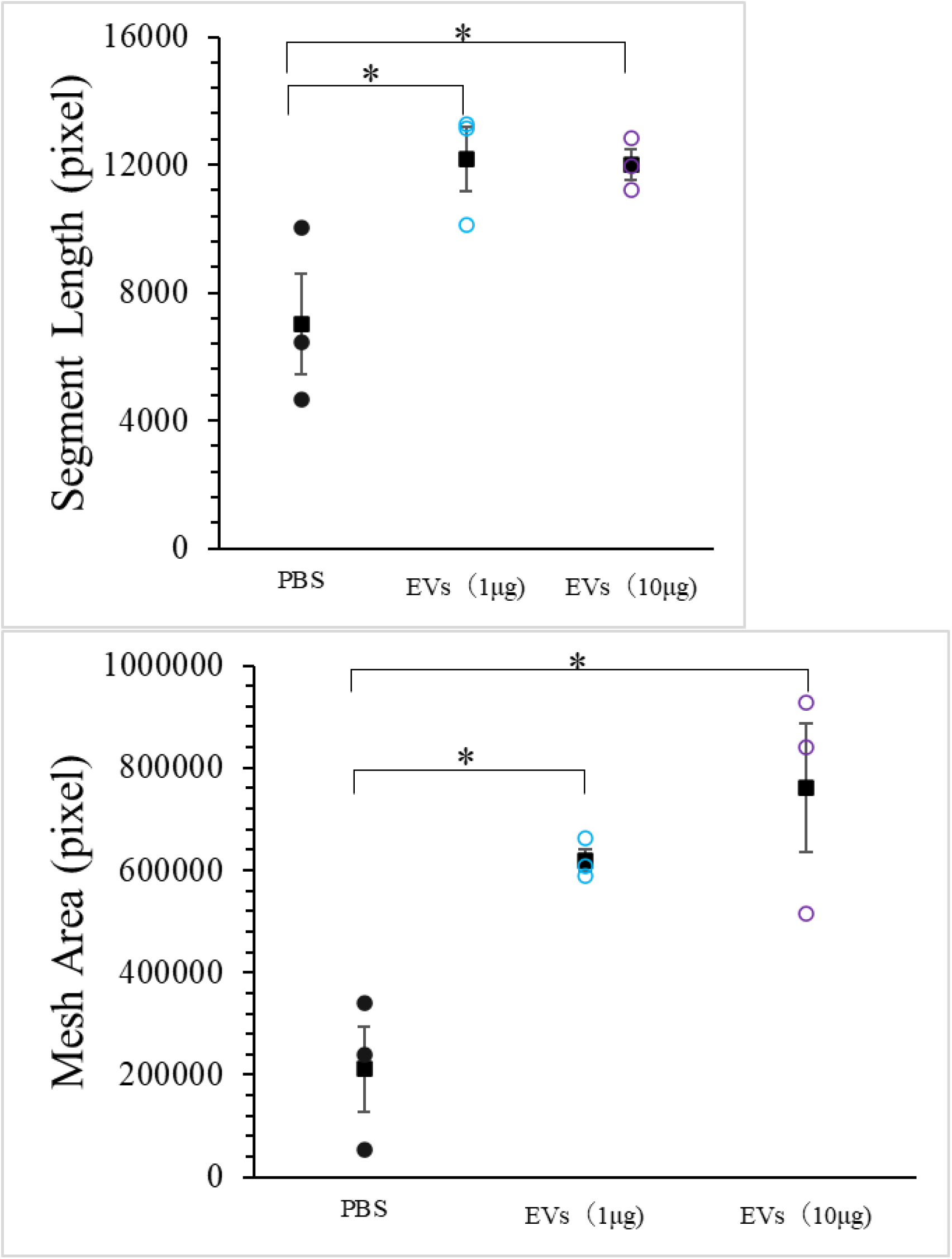
Analysis of tube formation in human umbilical vein endothelial cells (HUVECs) cultured with EVNative. A, HUVECs were cultured with EVNative in which protein concentration was diluted to 1 μg/ml. B, HUVECs were cultured with EVNative in which protein concentration was diluted to 10 μg/ml. C, HUVECs cultured with phosphate-buffered saline. D, Tube formation parameters were compared between the groups. N = 3 samples for each group. Data are presented as the mean ± standard error of the mean. *P<0.05 (one-way ANOVA).

**Figure 5.**
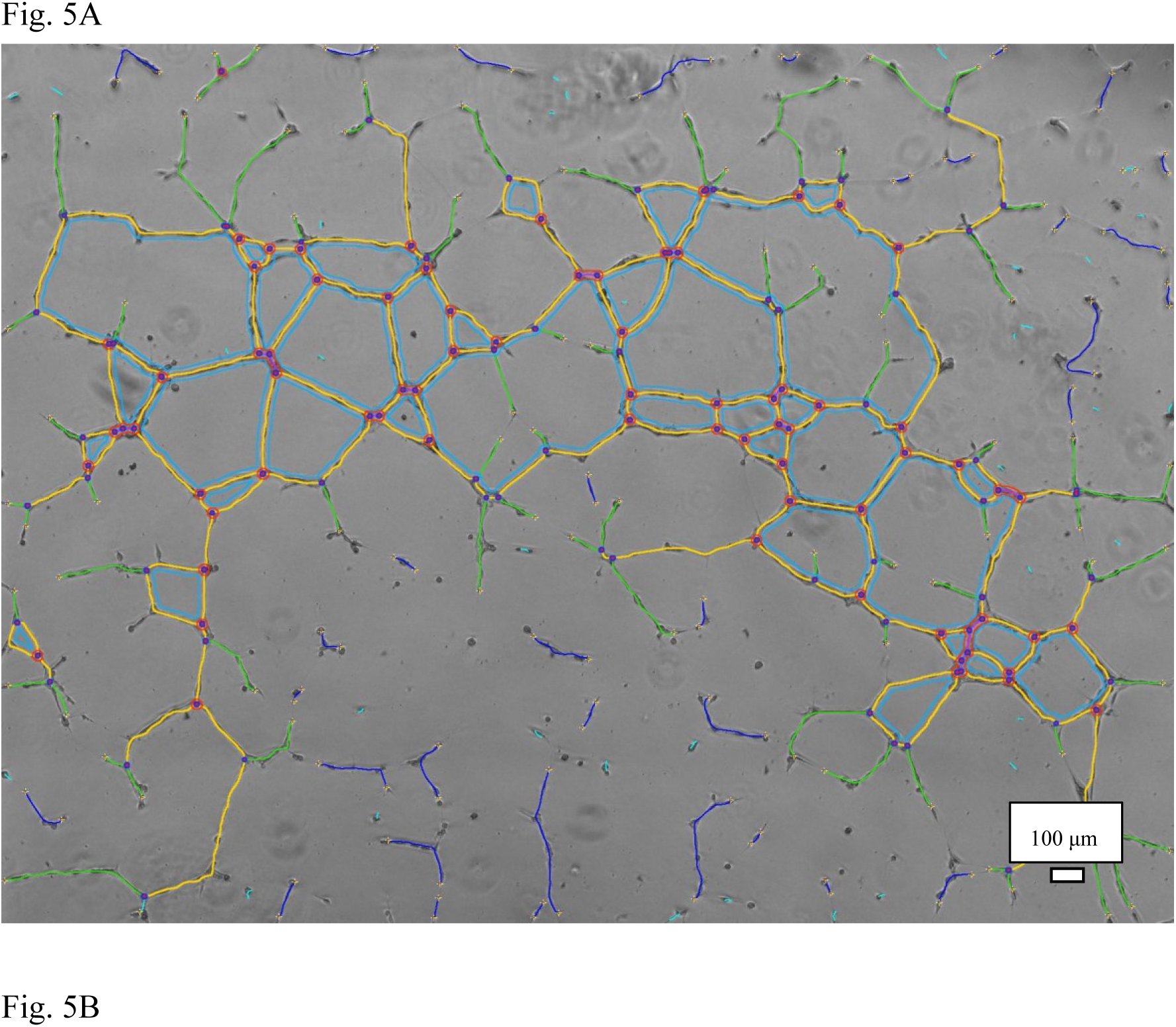
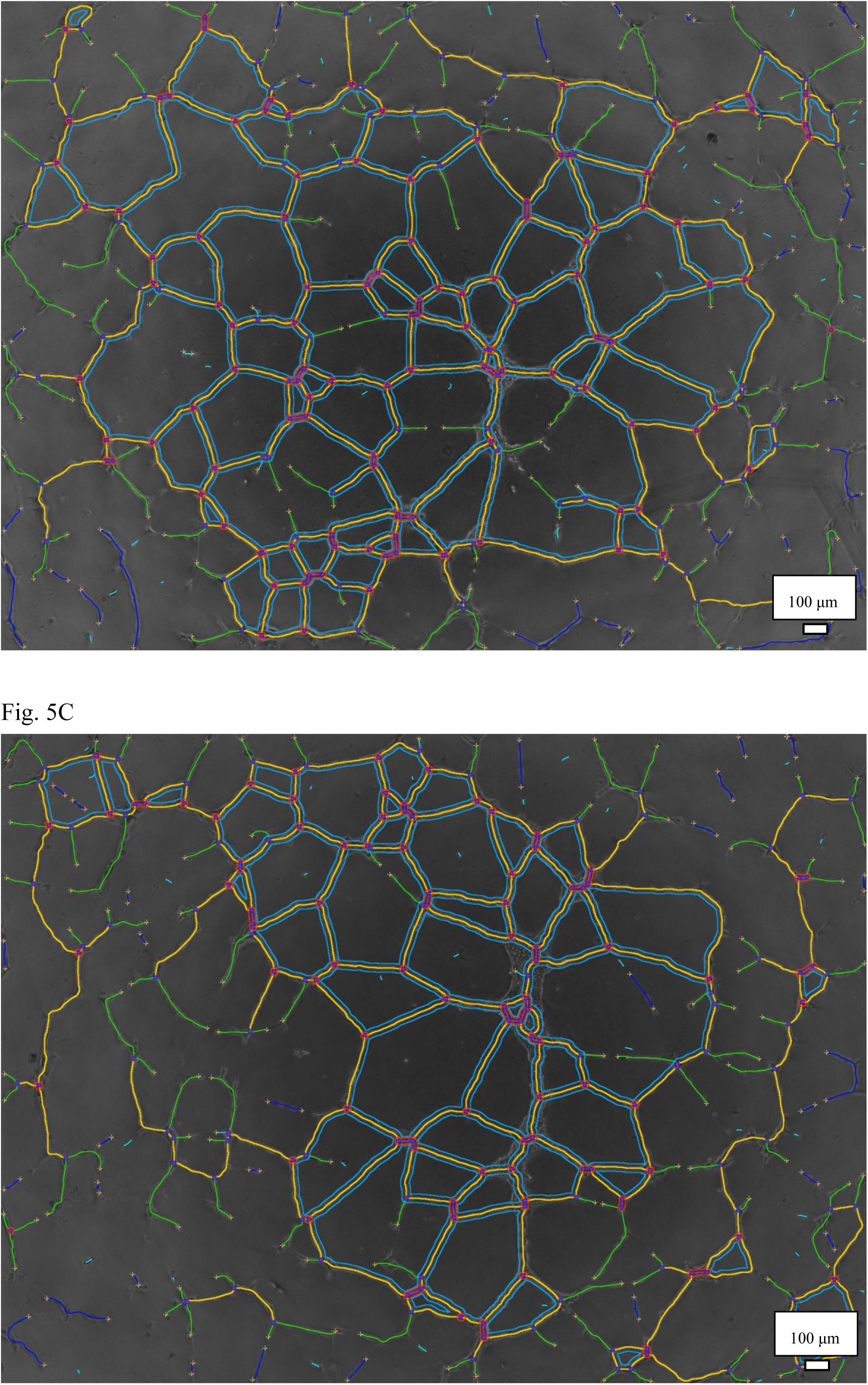
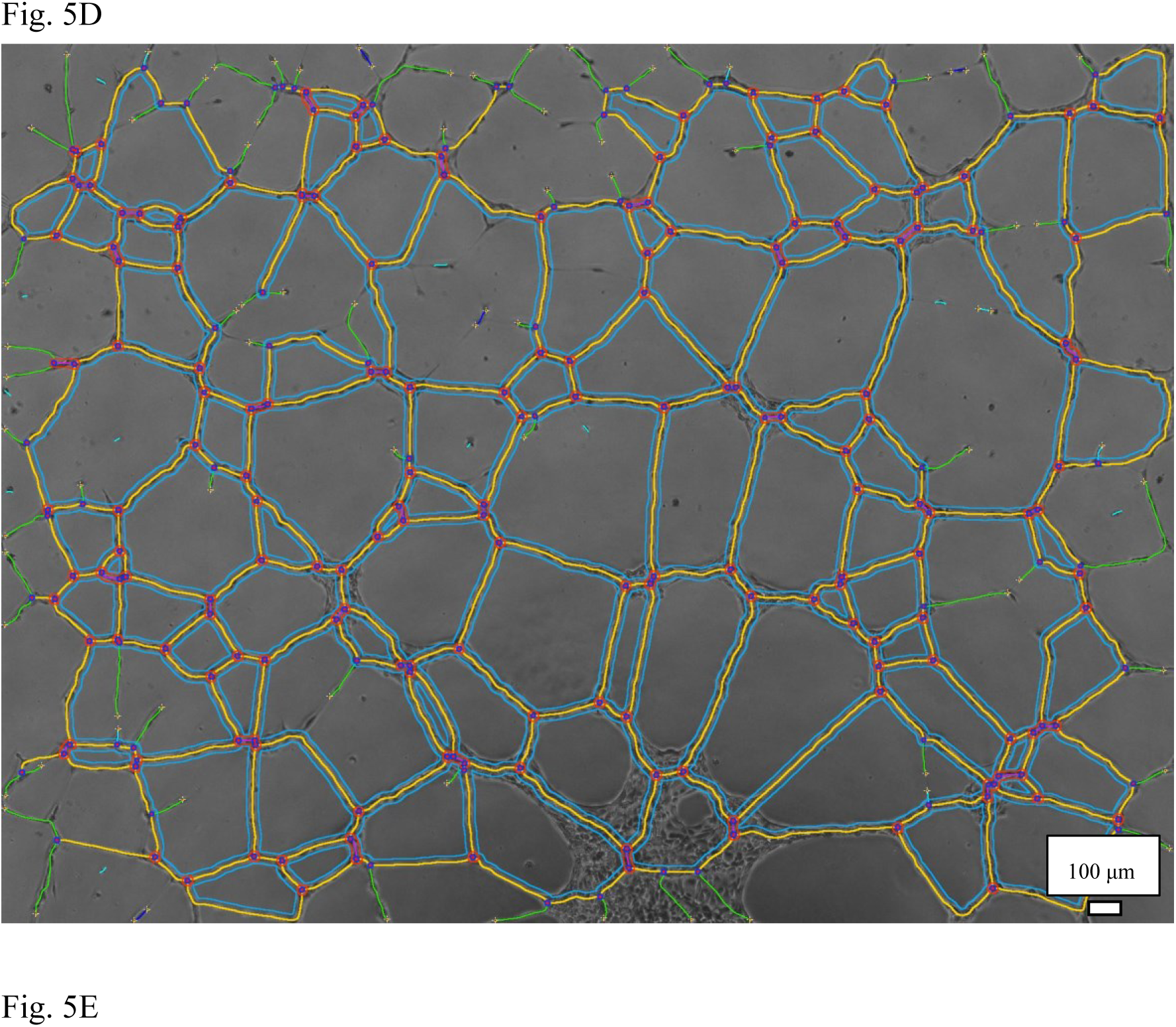
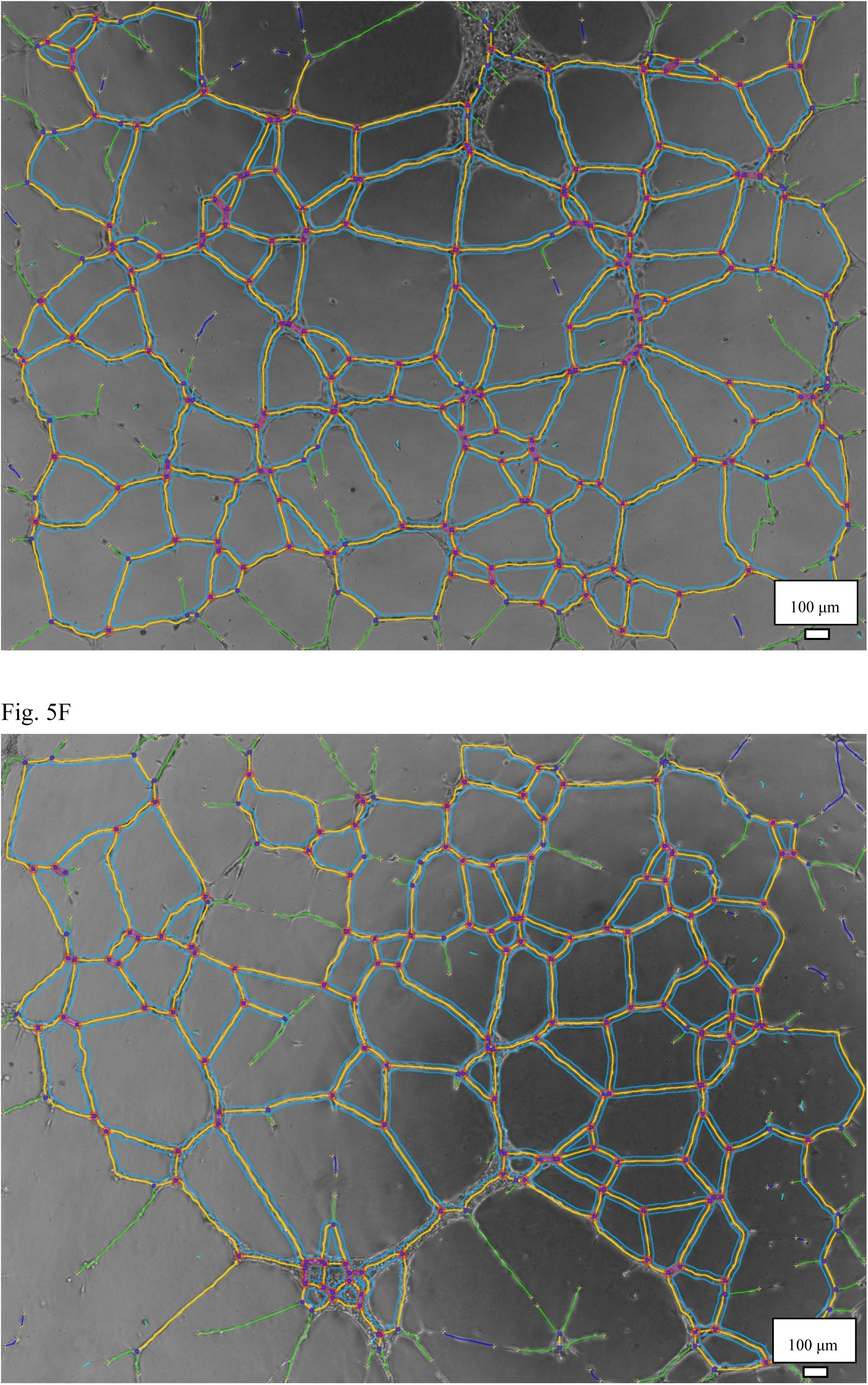
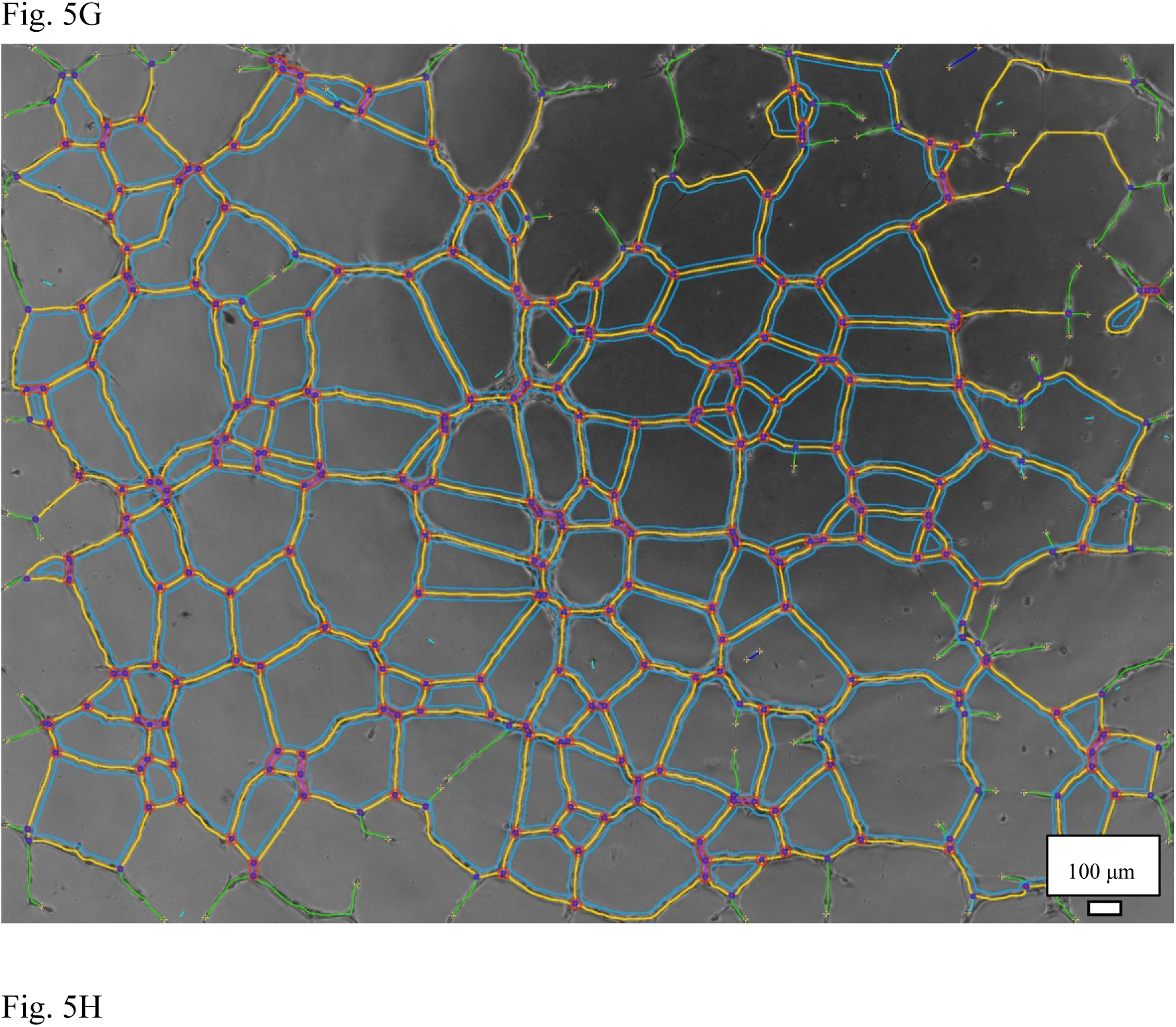
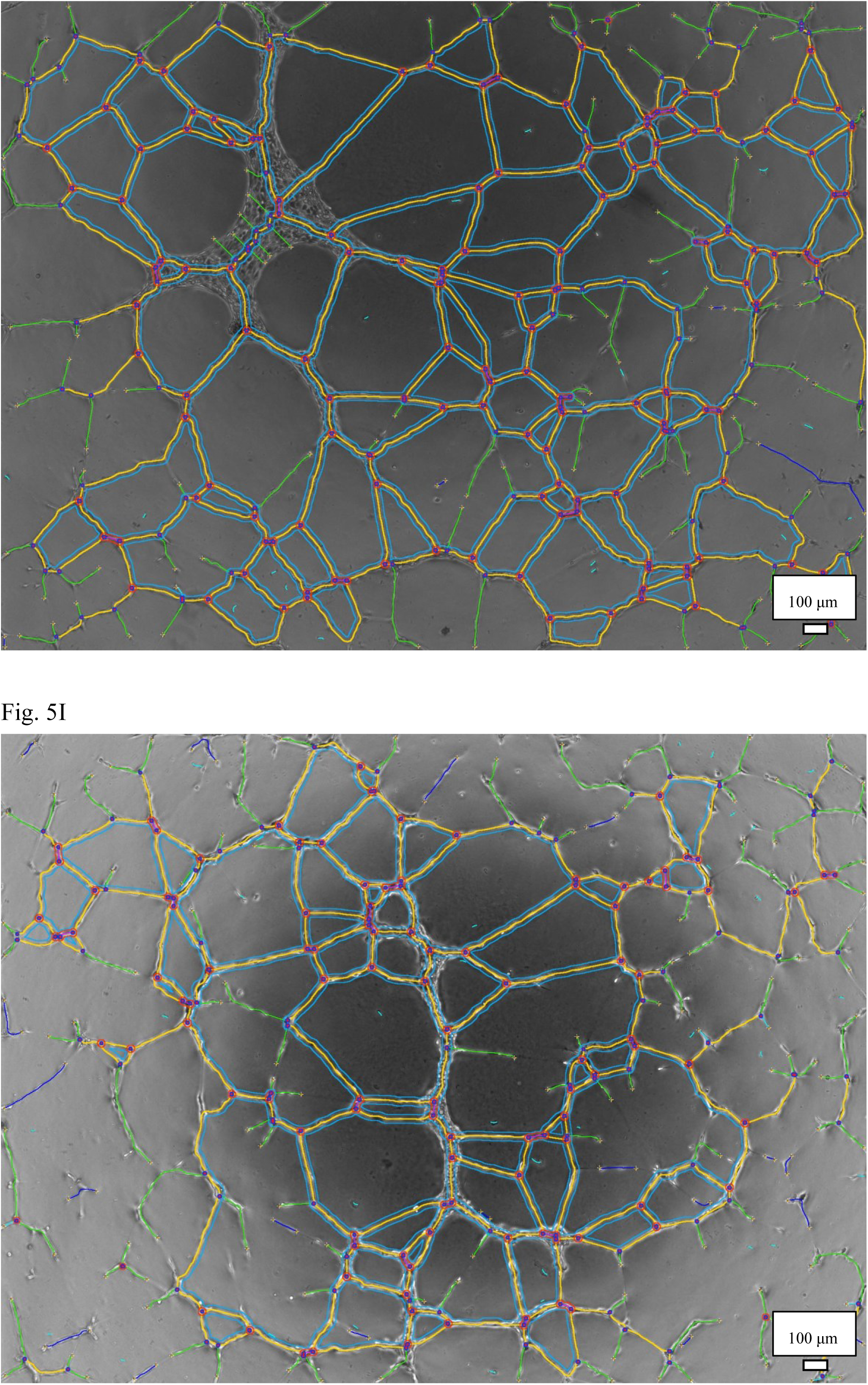
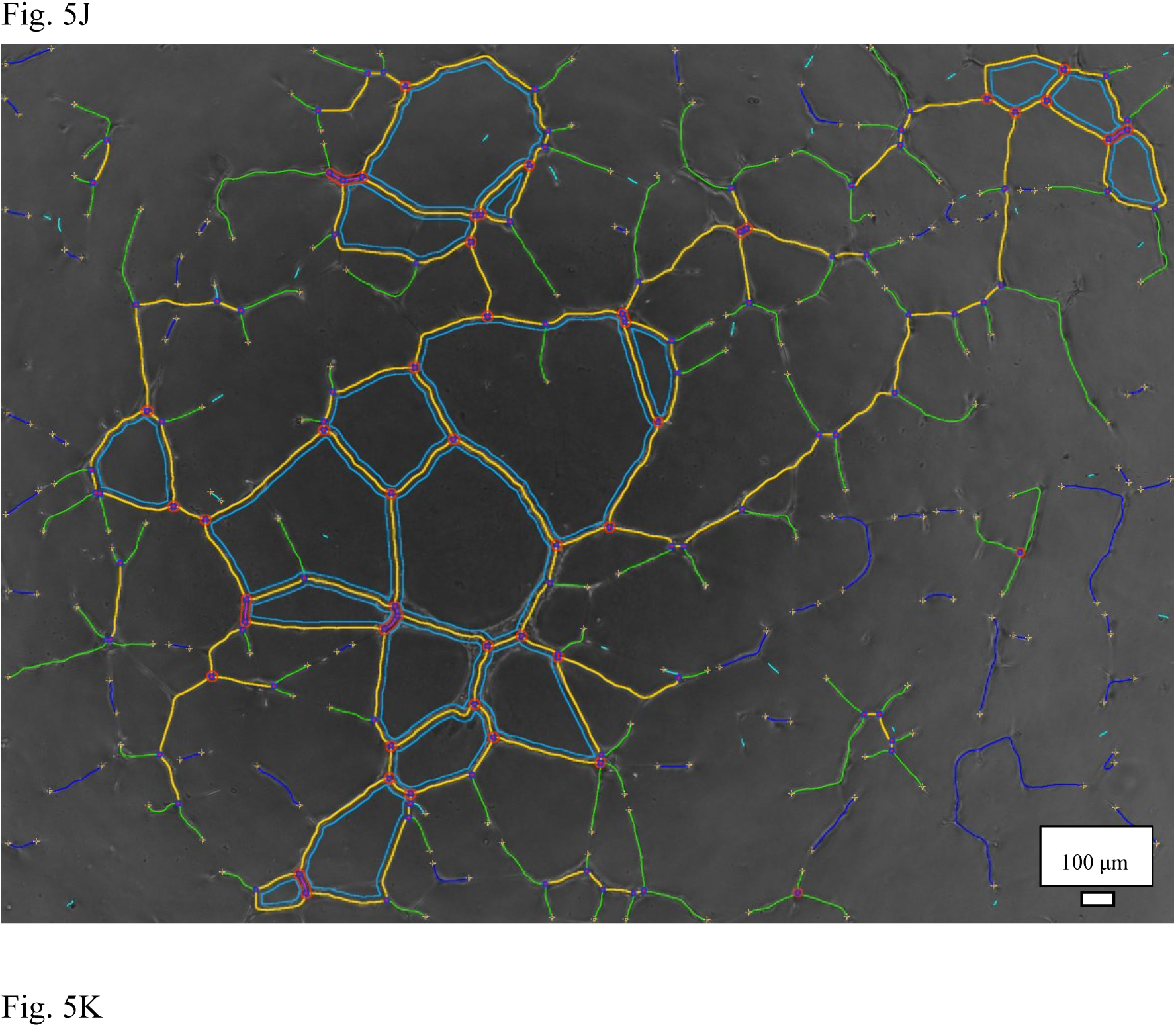
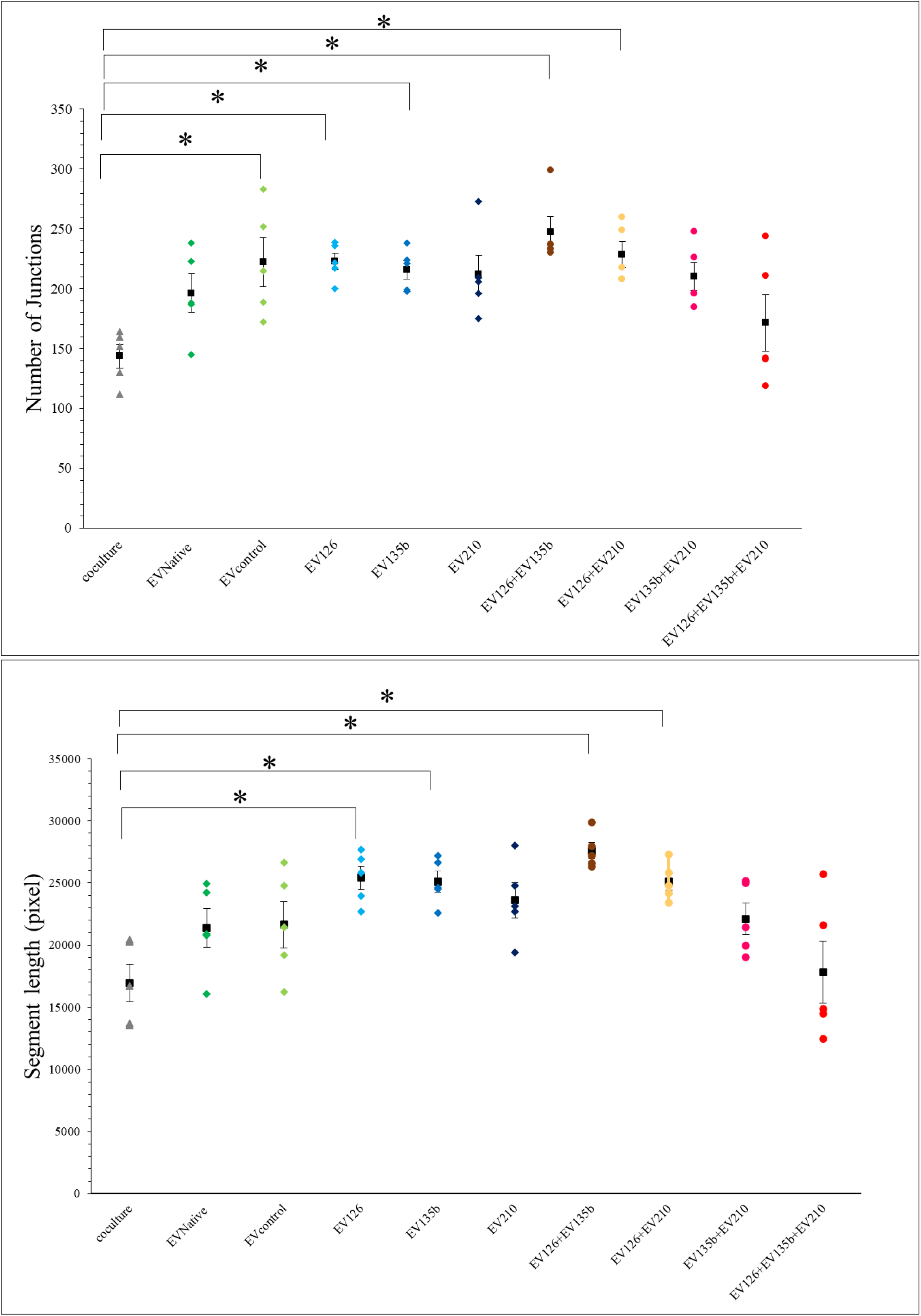
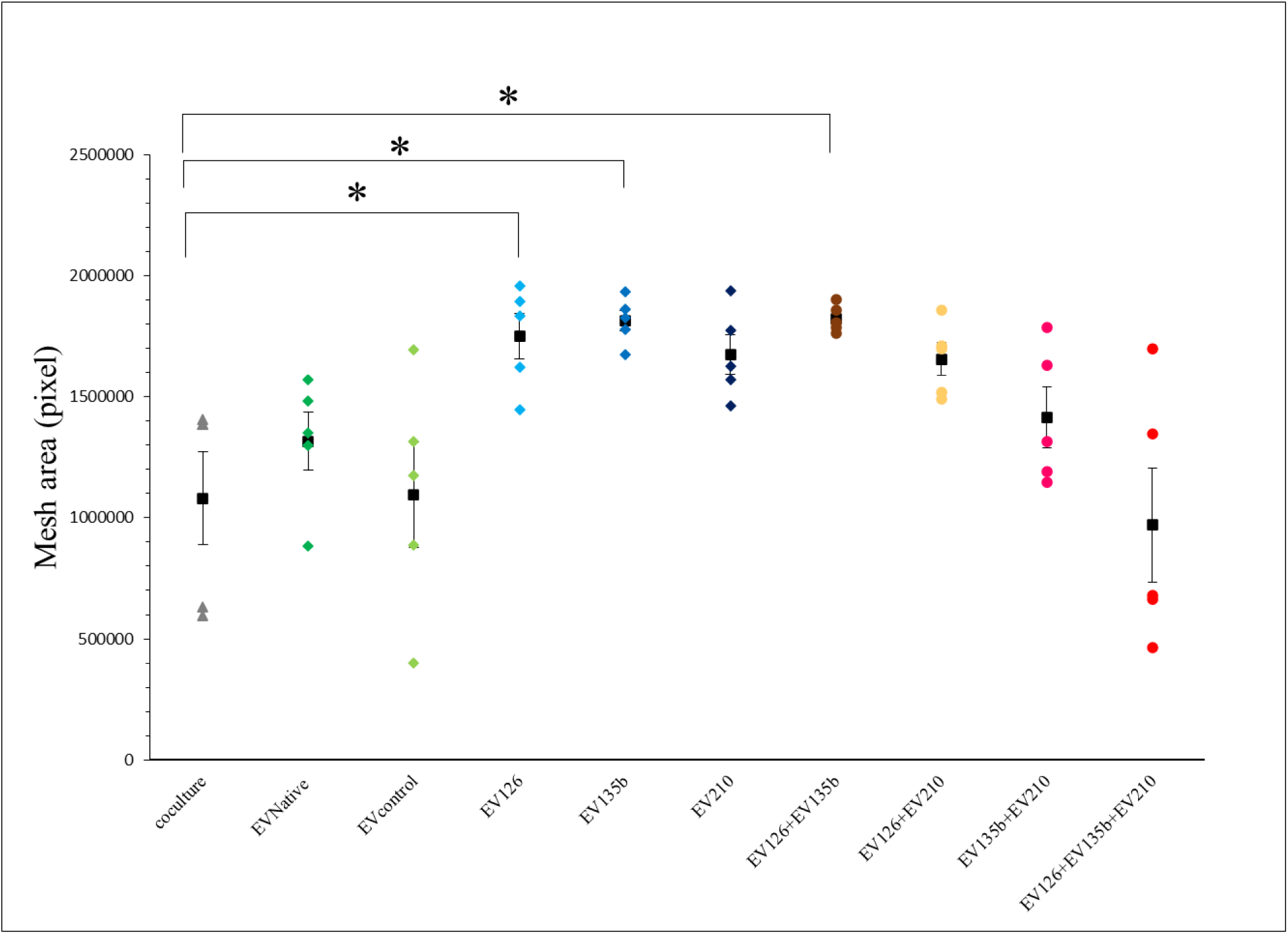
Analysis of tube formation in human umbilical vein endothelial cells (HUVECs) co-cultured with bone marrow-derived mesenchymal stromal cells (BM-MSCs) and extracellular vesicles (EVs). A, HUVECs co-cultured with BM-MSCs. B, HUVECs cultured with EVNative. C, HUVECs cultured with EVcontrol. D, HUVECs cultured with EV126. E, HUVECs cultured with EV135b. F, HUVECs cultured with EV210. G, HUVECs cultured with EV126 and EV135b. H, HUVECs cultured with EV126 and EV210. I, HUVECs cultured with EV135b and EV210. J, HUVECs cultured with EV126, EV135b, and EV210. K, Comparison of tube formation parameters between the groups. N = 5 samples in each group. Data are presented as the mean ± standard error of the mean. *P<0.05 (one-way ANOVA).

### Hindlimb ischemic model mouse establishment

Our results showed that 97.7% (43 /44) of mice exhibited loss of toe perfusion after femoral artery ligation and excision (Figure 6A-6B). Analysis of limb necrosis grading on day 7 demonstrated that mice injected with the combined EVs had lower necrosis grades than mice injected with PBS(–), EVNative, or EVcontrol (Figure 6C-6E). Blood flow analysis on day 7 revealed no significant differences in limb perfusion between mice injected with EVNative and EVcontrol. Furthermore, limb perfusion in the double EVs and triple EVs groups recovered significantly more than that in the EVcontrol group (P < 0.05) (Figure 6F).

**Figure 6.**
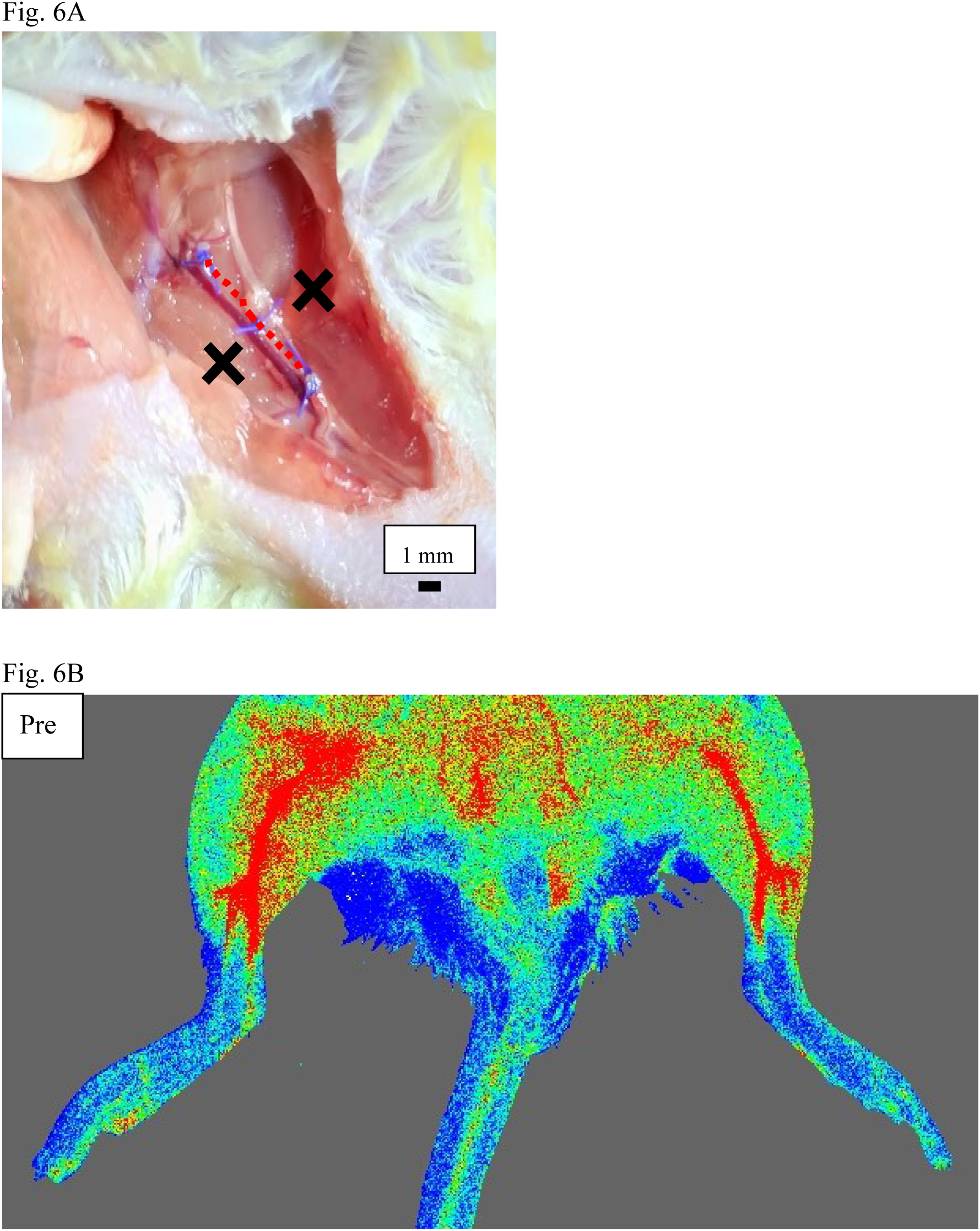
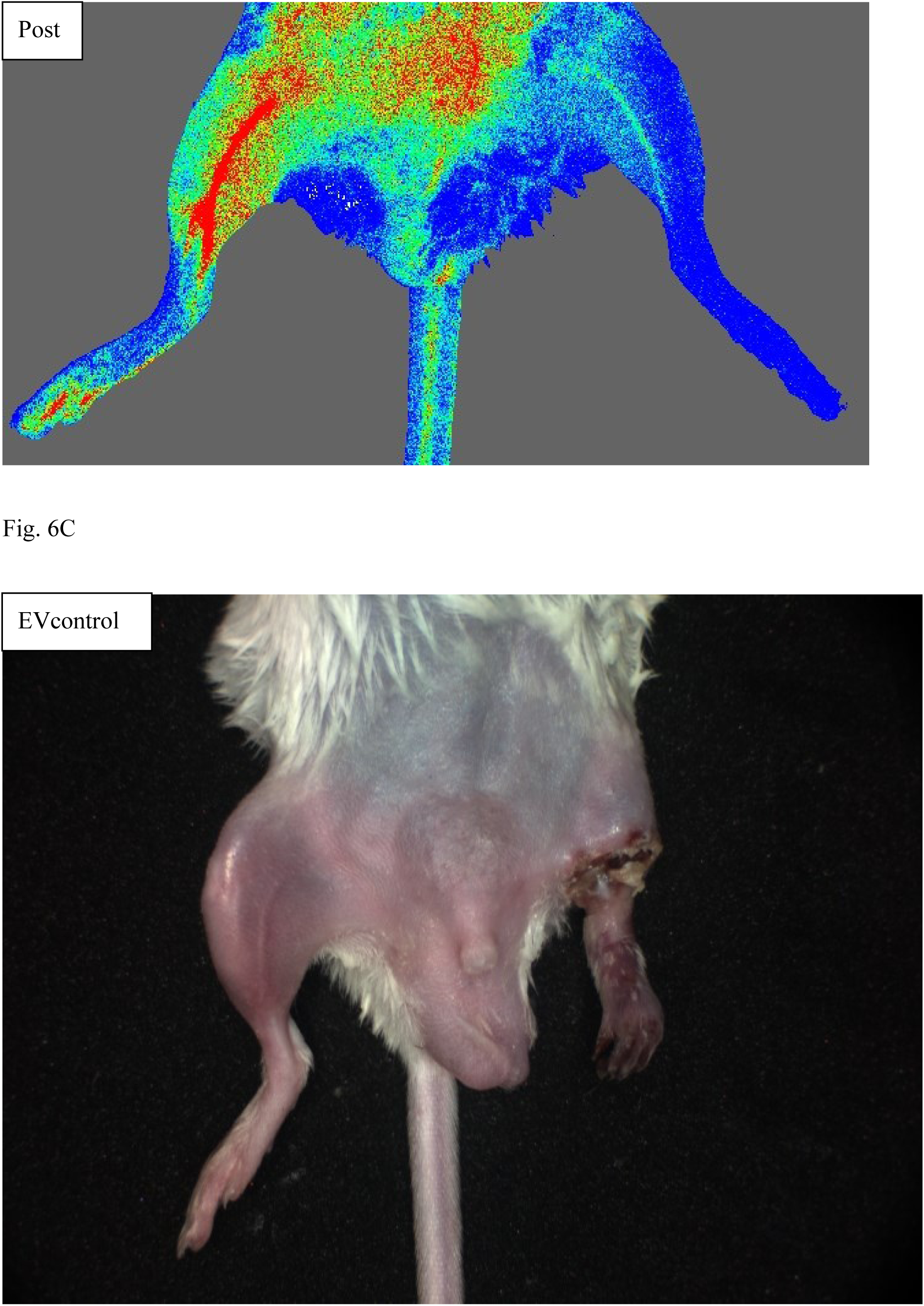
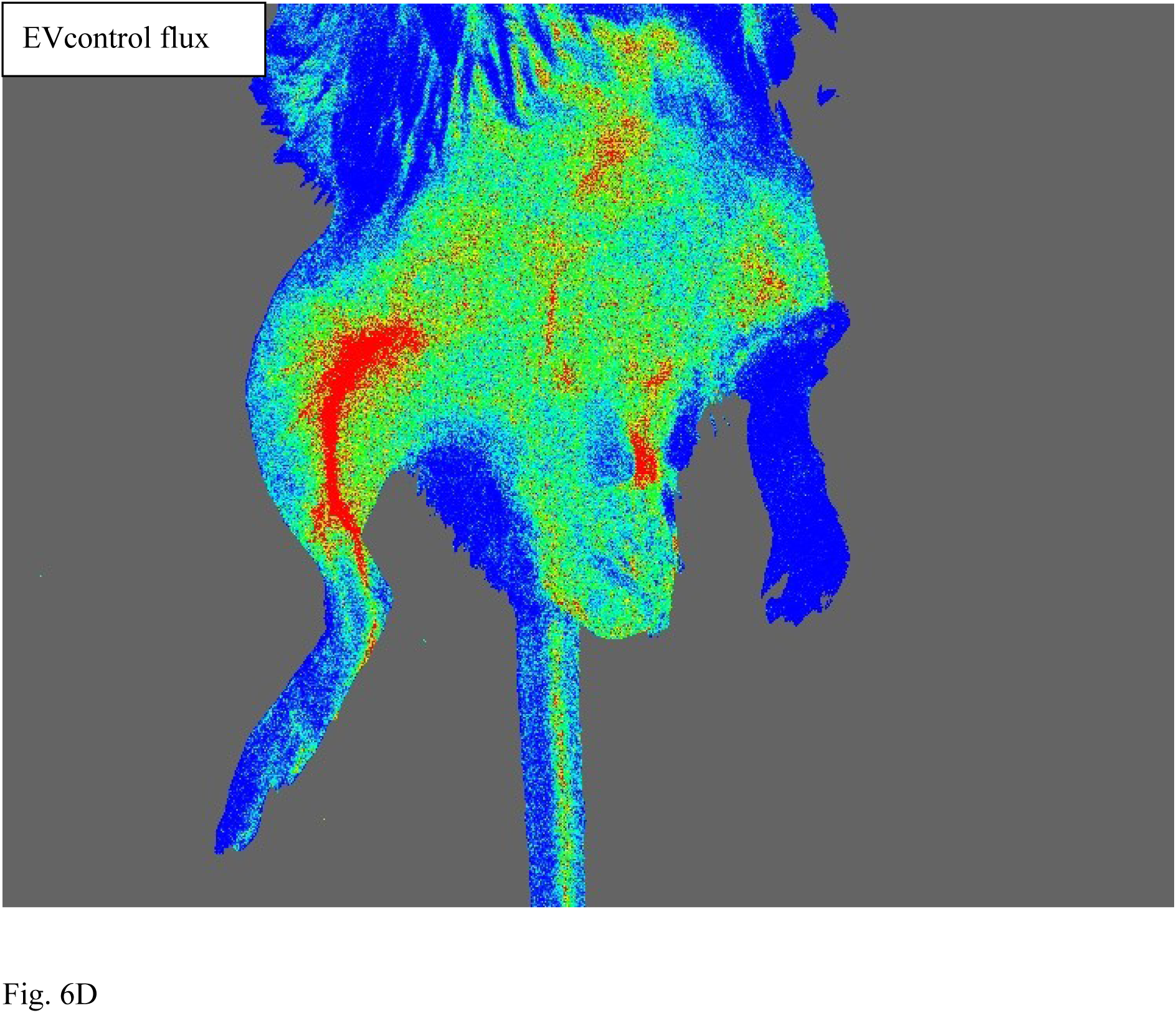
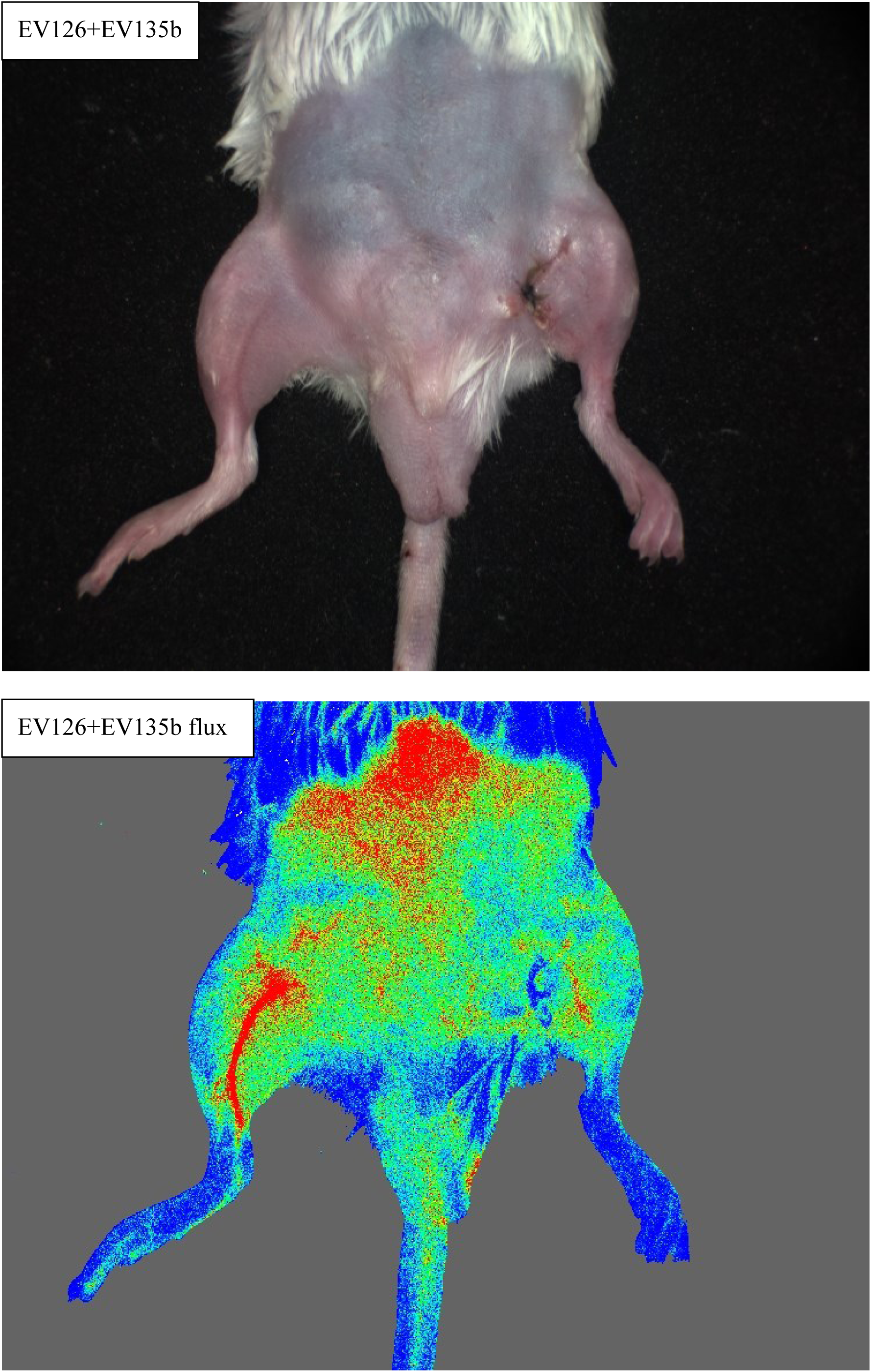
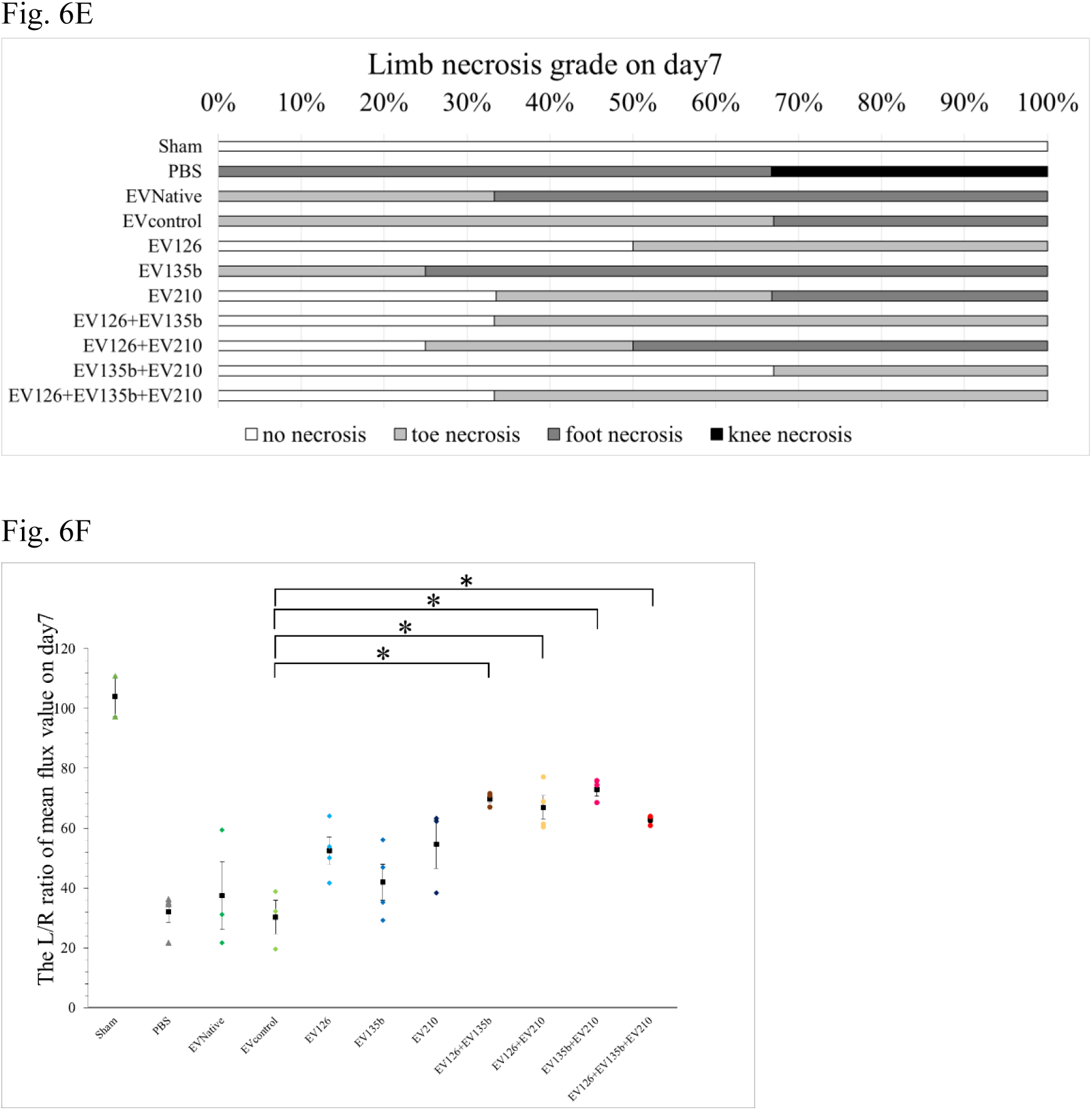
Mouse hindlimb ischemia model. A, Ligation and dissection of the left femoral artery. The red dotted line indicates the location of the femoral artery. Black crosses represent injection sites. B, Measurement of limb perfusion pre- and post-procedure using Moor FLPI-2. C, Respresentative hindlimb in an ischemic mouse injected with EVcontrol showed foot necrosis. D, Representative hindlimb in an ischemic mouse injected with combined EVs (EV126 and EV135b) showed no necrosis. E, Grading of the severity of limb necrosis in each group. The data showed that the mice injected with the combined EVs had a lower necrosis grade than the other groups. F, Left-right ratio of the mean flux value on day 7. Limb perfusion in the combined EVs group recovered significantly more than in the PBS(–) or single EVs groups.

Immunohistochemical staining of muscle tissue revealed that the number of the newly formed capillaries increased in the muscles injected with combined EVs compared with those injected with EVcontrol (Figure 7A). Histological analysis showed that the ratio of capillary endothelial cells to the ischemic muscle area was higher in mice injected with combined EVs than in those injected with EVcontrol (Figure 7B).

**Figure 7.**
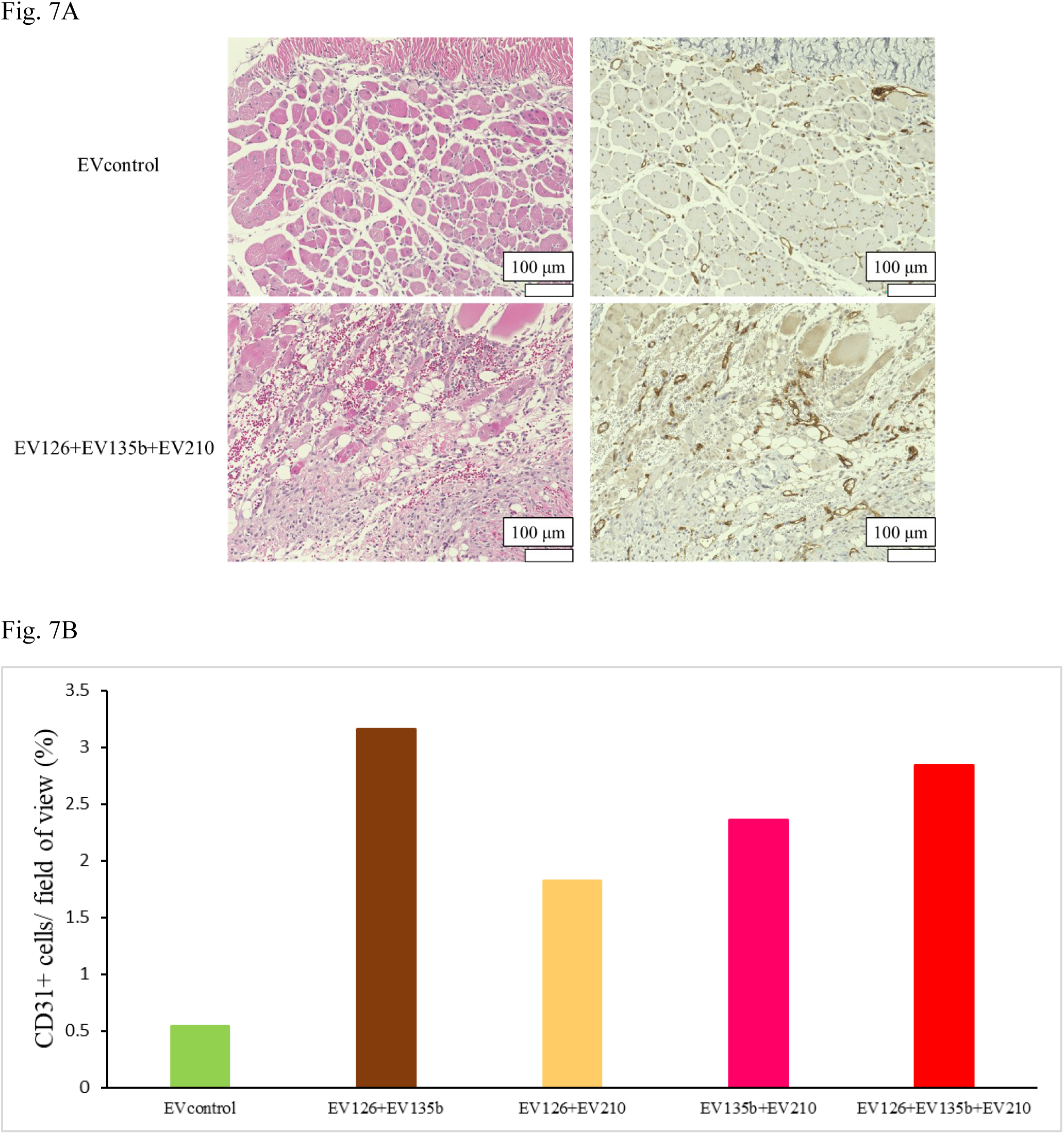
Histological analysis of the gastrocnemius muscles of ischemic limbs (n=1). A, Representative image of hematoxylin and eosin staining and immunohistochemical staining for CD31. B, Ratio of the area occupied by capillary endothelial cells in the ischemic muscle between mice injected with the EVcontrol and combined EVs.

## Discussion

We demonstrated that specific combinations of angio-miRNAs enhanced the angiogenic potency of EVs both *in vitro* and *in vivo*. To our knowledge, this is the first study to evaluate the angiogenic potency of EVs using specific combinations of angio-miRNAs.

EV heterogeneity poses a challenge in the accurate evaluation of studies utilizing EVs. In this study, we adhered to the MISEV2023 as much as possible to separate and use standardized EVs. In MSC-miRNA135b, the number of cells used to separate EVs was lower than that in the other groups, owing to their slow proliferation. Typically, this reflects lower protein and particle concentrations; however, the results of particle size distribution and the protein-to-particle ratio were similar to those of the other groups. This suggests that although the total number of collected EV135b particles decreased, the characteristics of the individual particles did not change.

In this study, we focused on the angio-miRNAs present in EVs derived from BM-MSCs. We successfully isolated EVs enriched with angio-miRNAs using lentiviral transfection. We found that lentiviral transfection did not affect BM-MSC proliferation or morphology; the fluorometry and NanoSight results confirmed this. Collectively, these findings suggest that lentiviral vector transfection is an effective method to produce functionally enhanced EVs without compromising their fundamental properties.

EVs derived from genetically modified BM-MSCs have superior angiogenic potency compared to those derived from the co-culture of BM-MSCs *in vitro*. Fukuda et al. conducted a clinical trial in which autologous BM-MSCs were injected intramuscularly into the limbs of patients with critical limb ischemia. This trial verified the safety of BM-MSC-based therapy and improved the limb amputation-free survival rate of patients^15^. However, while BM-MSCs have angiogenic potency, only a few differentiate into endothelial cells^16^. Instead, they promote angiogenesis in other cells through the paracrine effect^17^. In our study, a co-culture of BM-MSCs and HUVECs was used as a model of BM-MSC injection therapy in a clinical trial. Our results suggest that EVs from genetically modified BM-MSCs may be a more potent resource than BM-MSCs for therapeutic angiogenesis in patients with CLTI.

We also demonstrated that the combination of EVs derived from genetically modified BM-MSCs had a higher angiogenic potency than EVcontrol *in vivo*. EVs secreted from MSCs possess angiogenic potency and are expected to be used in regenerative medicine^18^. Several attempts have been made to modify EVs to enhance their angiogenic potency. Previous studies have shown that lentiviral vector transfection resulted in the overexpression of specific miRNAs in BM-MSCs and that EVs collected from BM-MSCs have enhanced angiogenic potency^19^. From our study, we hypothesized that combining several types of EVs containing angio-miRNAs could enhance angiogenic potency to a greater extent than individual EVs. Our preliminary research indicated that BM-MSCs expressed miRNA-126, -135b, and -210 (data not shown), all of which are known angio-miRNAs. We then produced three types of genetically modified BM-MSCs overexpressing each angio-miRNA and tested whether combinations of EVs from genetically modified BM-MSCs could promote angiogenesis. Although we successfully demonstrated that the combined EVs enhanced angiogenesis, it was remarkable that the observed angiogenic effect was not additive. The addition of double EVs to HUVECs did not result in double tube formation, and the injection of triple EVs into the ischemic hind limb of mice did not triple the recovery of limb perfusion. We speculated that they were related to the interactions between angio-miRNAs. Although these three angio-miRNAs have angiogenic potency, they act through different pathways. For instance, miRNA-126 regulates the response of endothelial cells to vascular endothelial growth factor (VEGF) by repressing phosphoinositol-3 kinase regulatory subunit 2 and sprouty-related *Drosophila* enabled/vasodilator-stimulated phosphoprotein homology 1 domain-containing protein 1, both of which are negative regulators of VEGF^20^. Moreover, miRNA-135 promotes angiogenesis by inhibiting hypoxia-inducible factor^21^, while miRNA-210 contributes to angiogenesis by inhibiting Ephrin A3^22^. The complex interactions of these angiogenic factors may make it difficult for combined EV injections to act additively. In addition, EVs injected into ischemic limbs were exposed to hypoxic conditions. Hypoxic conditions and ischemic tissue may affect the angiogenic reaction of EVs, especially miRNA-135b and -210; this was reflected in the different angiogenic results between our *in vitro* and *in vivo* experiments.

In this study, EVs were injected into thigh muscles from the surgical site. The injection site and timing were not strictly standardized in previous studies^23-26^. We injected EVs into thigh muscles from the surgical site to allow for direct observation and precise injection. The high rate of toe ischemia after femoral artery excision, as proven by our MoorFLPI-2 results, supports the accuracy of our hindlimb ischemia model. Percutaneous oxygen saturation was also measured; however, the pulse wave was too weak for oxygen saturation to be detected. Laser speckle imaging has been used to evaluate visceral perfusion^27-28^. Visual analysis of peripheral blood perfusion using MoorFLPI-2 enabled the precise evaluation of reperfusion after EV injection.

This study had two limitations. First, the evaluation period for the ischemic limbs was relatively short. Each group initially contained four mice, and muscle tissue was harvested on day 14 from one individual in each group. We planned to continue the evaluation of limb perfusion on days 21 and 28, but this was not possible because the mice already died due to limb gangrene. Second, we did not investigate changes in protein or gene expression during the angiogenic processes. Future studies should investigate how these EV miRNAs act when delivered to endothelial cells to elucidate the molecular mechanisms underlying our results.

## Conclusion

In this study, we demonstrated the angiogenic effects of a combination of three specifically modified EVs both *in vitro* and *in vivo*. Our study will contribute to clinically applicable use of modified EVs for regenerative therapy in CLTI.

## Non-standard Abbreviations and Acronyms

BM-MSC: bone marrow-derived mesenchymal stromal cell
CLTI: chronic limb-threatening ischemia
EVs: extracellular vesicles
HUVECs: human umbilical vein endothelial cells
miRNA: micro RNA
RFP: red fluorescent protein
VEGF: vascular endothelial growth factor

## Acknowledgements

We thank M. Abe and K. Sudo for technical support and A. Matsuda, I. Mizoguchi, and T. Yoshimoto for their scientific advice.

## Author Contributions

Y. Wada performed the experiments, analyzed the data, and wrote the article. S. Fukuda conceived the project, supported the surgical procedures, and edited the manuscript. T. Kusakabe and A. Inoue assisted with the cell culture and experiments. Y. Yoshioka provided technical support for the ExoScreen assay. T. Ochiya provided scientific advice and reviewed the article critically. T. Kudo supported the surgical procedure.

